# The natural compound Notopterol targets JAK2/3 to ameliorate macrophage-induced inflammation and arthritis

**DOI:** 10.1101/832782

**Authors:** Qiong Wang, Xin Zhou, Long Yang, Yongjian Zhao, Jun Xiao, Qi Shi, Qianqian Liang, Yongjun Wang, Hongyan Wang

**Affiliations:** Longhua Hospital, Shanghai University of Traditional Chinese Medicine, 725 Wan-Ping South Road, Shanghai 200032, China; State Key Laboratory of Cell Biology, Key Laboratory of Systems Biology, CAS Center for Excellence in Molecular Cell Science, Shanghai Institute of Biochemistry and Cell Biology, Chinese Academy of Sciences, University of Chinese Academy of Sciences, Innovation Center for Cell Signaling Network, Shanghai, 200031, China; Cancer Center, Shanghai Tenth People’s Hospital, Tongji University, School of Medicine, Shanghai 200072, China; Department of rehabilitation medicine, Shanghai Eighth People’s Hospital

**Keywords:** Notopterol, JAK2/3, macrophage, inflammation, collagen-induced arthritis

## Abstract

Notopterol (NOT) is one of the main constituents of the traditional Chinese medicinal herb *Notopterygium incisum Ting ex H.T. Chang* has anti-rheumatism activity, but the target of NOT remains unknown. Here we have demonstrated that orally or intraperitoneal administration of NOT exhibits significant therapeutic effects on the collagen-induced arthritis (CIA) model in both DBA/1J and C57/BL6 mice. NOT treatment *in vivo* and *in vitro* reduces production of inflammatory cytokines and chemokines in TNFα- or LPS/IFNγ-stimulated macrophages via blocking the JAK2/3-STAT3/5 activation. Mechanistically, NOT directly binds JAK2 to inhibit its activity via Arg980, Asn981, and Leu932 in the JH1 domain. Importantly, expression of the L938A/R980A/N981A mutant in zebrafish significantly inhibited the *in vivo* inflammatory response after LPS injection, which showed no further inhibitory effect upon NOT treatment. Combination of NOT and an anti-TNFα antibodies could achieve a better therapeutic effect than anti-TNFα alone in the CIA model. We therefore suggest that as a specific JAK2/3 inhibitor, the natural compound NOT ameliorates pathology of RA, which might be useful to treat other JAK2/3-related diseases.

## Introduction

Rheumatoid arthritis (RA) is a chronic and systemic autoimmune disorder that primarily affects the joints, resulting in pain, swelling and irreversible deformities ^1^. Various studies have suggested that macrophages participate in the pathogenesis of RA by producing multiple cytokines for a long term to cause tissue damage ^2, 3, 4, 5, 6^. In addition, the TLR4 pathway induced cytokines were reported that highly related to RA patients ^2, 3^. Several biologic agents against specific cytokines and their receptors have led to substantial progress in RA treatment, such as anti-antibodies to tumor necrosis factor (TNF) and IL-6 ^7, 8^. However, these biologicals increase the risk of opportunistic infections, and are limited in their broad use because of high costs and the need to be injected. Thus, a need still exists for orally available, small molecules that are effective in the therapeutic treatment of RA^9, 10^.

Activation of the JAK/STAT pathway (JAK, janus kinases; STAT, signal transducers and activators of transcription) induces the expression of multiple inflammatory cytokines and chemokines in macrophages^11, 12^. In particular, the autocrine loop between TNF/IFN and JAK/STAT could trigger and amplify the expression of multiple inflammatory cytokines ^13, 14^. Upon cytokine stimulation, the relevant receptors recruit JAK via their intracellular domains, resulting in phosphorylation of JAK at tyrosine residuesand then the recruitment and phosphorylation of STAT. The phosphorylated STATs, in turn, form homo- and heterodimers, translocate rapidly to the nucleus, and activate a suite of gene transcription. In mammals, there are 4 members of the JAK family (JAK1, JAK2, JAK3 and Tyk2) and 7 STATs (STAT1/2/3/4/5a/5b/6), resulting in a complex downstream signaling pathway. Because the JAK/STAT pathway plays an important role in RA ^15^, blocking its signaling suppresses the inflammatory response in macrophages to achieve a therapeutic effect on RA progression^9, 12^. For example, Tofacitinib, the first-generation inhibitor of the JAK family (JAK1 and JAK3 especially), has been approved by the US FDA for the treatment of RA in patients with intolerance or non-response to Methotrexate ^11, 14, 16, 17, 18, 19^.

*Notopterygium incisum Ting ex H.T. Chang*, which is a traditional Chinese medicinal herb, which has been used for the treatment of rheumatism ^20, 21, 22, 23, 24^. Nevertheless, given its complex composition and unclear pharmacology, its further clinical application to treat RA worldwide or treat other clinical diseases is limited. Notopterol (NOT) is one of the main constituents of this herb and has been reported to have analgesic properties, including the inhibition of acetic acid-induced writhing in mice, as well as displaying anti-inflammatory activity as evidenced by an inhibitory effect in the vascular permeability test^25, 26^. However, the direct molecular target(s) of NOT and whether it can ameliorate RA pathology are unknown. In this study, we demonstrated that NOT dampens inflammation and ameliorates arthritis on several mouse CIA models while achieving an improved therapeutic effect when combined with an anti-TNFα antibody. In addition, mechanistaltly, we have elucidated that NOT directly binds the JH1 domain of JAK2 to inhibit JAK2 activation leading to reduction of inflammatory cytokine or chemokine production in macrophages without affecting macrophage apoptosis. It indicated that NOT has potential therapeutic effect on other macrophage associated-diseases.

## Experimental procedures

### Mice and ethics statement

6-8 weeks old, male or female, C57/BL6 and DBA/1J mice were obtained from Shanghai Slack Laboratory Animals Co., Ltd, and used for the induction of CIA model. The animal studies were approved by the Institutional Animal Care and Use Committee (IACUC) of Institute of Biochemistry and Cell Biology, Shanghai Institutes for Biological Sciences, Chinese Academy of Sciences officially (approval numbers 1608-023).

### Culture of primary macrophages and cell lines

Raw 264.7 and 293T cells were obtained from ATCC (ATCC® Number was CRL-11268™ and TIB-71™ respectively). Bone marrow-derived macrophages (BMDMs), peritoneal macrophages (PEMs), 293T, and RAW264.7 were cultured at 37 °C, 5% CO2 in complete DMEM medium (Invitrogen, Grand Island, CA, USA) consisting of 10% (vol/vol) FBS and 100 U/ml P/S.

To obtain BMDMs, the lower limbs of C57/BL6 mice were cut off, muscle around the tibia and femur was removed, the ends of the bones were subtracted. Five-milliliter syringe full of buffer consisting of DMEM, 100U/ml penicillin (Invitrogen), 100U/ml streptomycin (Invitrogen), 1% L-Glutamine (Invitrogen), 1%FBS (Invitrogen), was used to wash the marrow cavity 3 to 5 times to obtain bone marrow cells. After filtered through 70-µm cell strainers and then centrifuged for 5 min at 1000 rpm, bone marrow cells were resuspended in 5 ml erythrocyte lysis buffer (154 mM NH4Cl, 10 mM KHCO3, 0.1 mM EDTA, pH 7.4). After centrifugation and washing, the cells were resuspended and cultured in macrophage-conditioned medium containing complete 1640 medium containing 10% (vol/vol) FBS and 100 U/ml penicillin and streptomycin and 20 ng/ml murine M-CSF (Peprotech, cat. # 315-02) for 7 days.

To get PEMs, 3 ml of 3% bacteria-free thioglycollate solution (Sigma, B2551-500G) was injected into the abdominal cavity of C57/BL6 mice. Four days later, 5 ml buffer was injected into the abdominal cavity, massaged for 2 minutes and collected the buffer. After centrifugation at 1000 rpm for 5 min, the cells were resuspended in 5 ml erythrocyte lysis buffer and filtered through 70-µm cell strainers. After centrifugation and washing, the PEMs were overnight incubated with DMEM complete medium for the following usage.

### Type II collagen induced arthritis (CIA) model and Notopterol treatment

Twice immunization to induce CIA : First immunization (D0): the emulsion (100 μL) containing completely emulsified chicken type II collagen (2 mg/ml, Chondrex, cat: # 20011) and complete Freund’s adjuvant (4mg/ml *M. tuberculosis,* Chondrex, cat: # 7001) at equal volume, was intradermal injected into the skin around the base of the tail. Secondary immunization (D21): the emulsion (100μL) containing completely emulsified chicken type II collagen and incomplete Freund’s adjuvant (Chondrex, cat: # 7002) at equal volume was intradermal injected next to the first injection site. The limbs showed obvious redness and swelling within one week and the incidence was greater than 90% for DBA/1J mice and 70% for C57BL6 mice.

One immunization to induce CIA : Bovine type II collagen (2 mg/ml, Chondrex, cat. # 20021) and complete Freund’s adjuvant (4mg/ml M. tuberculosis, Chondrex, cat. # 7001) were completely emulsified and subcutaneous injected into the tail (100 μL each). The limbs displayed obvious redness and swelling from Day 42 to Day56, and the incidence was above 90%.

Notopterol treatment: NOT (ChemFaces Biochemical Co., Ltd., cat. # CFN98563, CAS: 88206-46-6) was dissolved in DMSO at 200 mg/ml and stored at −80°C in the dark. Before injection, NOT was diluted 200 times with saline (1 mg/ml). For the intragastric administration, NOT (1 mg/kg) was suspended in 0.5% sodium carboxyl methyl cellulose (CMC-Na). For the CIA model with two immunization, NOT administration started at day 22. For the CIA model with one immunization, NOT treatment began at day 42.

### Clinical score

Clinical score of arthritis was evaluated: grade 0, normal and no swelling; grade 1, mild, redness and swelling of the ankle or wrist, or apparent redness and swelling limited to individual digits, regardless of the number of affected digits; grade 2, moderate redness and swelling of ankle or wrist; grade 3, severe redness and swelling of the entire paw including digits; grade 4, maximally inflamed limb with involvement of multiple joints^27, 28^. Each limb was graded with a score of 0–4, with a maximum score of 16 for each mouse and 0 for the minimum. Two independent examiners unaware of the group assessed clinical score.

### Histology, Immunofluorescence and tissue preparation

The ankle and knee joints were decalcified by 10% EDTA in PBS (PH7.4) before dehydration. The kidney, lung, spleen, and liver from mice were cut into 5 micrometers (μm) slices. The slices were stained by Alcian Blue/Orange G (ABOG), hematein eosin (H&E) or tartrate-resistant acid phosphatase (TRAP) staining. Synovium area, cartilage area and talar area at ankle joints and synovium area and cartilage area at knee joints were analysed by Adobe Acrobat Pro software. Images were acquired by Olympus BX-51 microscope. For immunofluorescence staining, trypsin-induced antigen retrieval was applied on rehydrated paraffin sections. After blocking with 5% BSA, primary antibodies including anti-F4/80 (Abcam, cat. # ab6640) and anti-iNOS (Abcam, cat. # ab15323) was applied to the sections overnight at 4 °C. After washing with PBS, a secondary antibody (Goat-anti-Rat IgG H&L, Alexa Fluor® 647, Abcam. cat. # ab150167 and Goat-Anti-Rabbit IgG H&L, Alexa Fluor® 488, Abcam. cat. # ab150077) was applied at the same time with DAPI (4’,6-diamidino-2-phenylindole, Sigma, cat. # D9564). The images were taken by FV1200 Laser Scanning Microscopes (Olympus). Representative images were selected in a blinded fashion.

Blood samples were obtained from the orbit and serum was precipitated after centrifugation. The hind paw (from the tibiotalar joint to the tip) was soaked in ice PBS and crushed mechanically to get paw tissue homogenate. The kits to measure aspartate aminotransferase (AST), alanine aminotransferase (ALT), creatinine (CRE) and urea nitrogen (UREA) were from Shanghai Shensuo UNF medical diagnostic articles co., LTD and the IL-1β, IL-6 and TNFα Elisa kits were from NeoBioScience.

### Micro computed Tomography (micro-CT) analysis

After fixing with 4% paraformaldehyde for 48 hours, the ankle joints were washed by PBS for 2 hours, soaked into 75% ethanol, and scanned by the micro-CT system (Scanco VIVA CT80, SCANCO Medical AG, Switzerland). The scanning parameters were pixel size: 15.6 μm, tube voltage: 55 kV, tube current: 72μA, integration time: 200 ms. The cross-section images were reconstructed and realigned in 3D, the bone volume of talus were measured and the density threshold was set from 370 to 1000 as “bone” by μCT Evaluation program V6.6 (Scanco Medical AG, Switzerland). A stack of 340-441 cross-sections was reconstructed, with an interslice distance of 1 pixel (15.6 μm), corresponding to a reconstructed height of 5.3-6.9 mm and recreating the ankle joints.

### The *in vivo* examination of inflammation using zebrafish

The mRNA of JH1 and JH1-3A was synthesized *in vitro* by mMESSAGE mMACHINE™ T7 Transcription Kit (Invitrogen™, AM1344). 100 pg (1nL) JH1 or JH1-3A were injected into one cell stage of zebrafish embryos (Tuebingen line) to generate zebrafish expressing JH1 or the JH1-3A mutant. Injection of LPS or LPS in combination with NOT was performed at 48 hours post-fertilization (hpf). Zebrafish mRNA was harvested at 54 hpf to measure the *in vivo* production of inflammatory cytokines.

### Quantitative RT–PCR and RNA-seq analysis

Total RNA was extracted from tissues or cells by using Trizol reagent (Invitrogen). Reverse transcription was performed to generate cDNA by using PrimeScriptTM RT reagent Kit (Takara, cat. # RR047A). Quantitative PCR was performed on LC480 (Roche) and SYBR Green Mix was from Shanghai Yeasen Biological Technology Co., Ltd. The relative mRNA levels, normalized to the expression of β-actin, were calculated by the ΔΔCt method. Primer sequences were listed in Table 1.

**Table 1.**
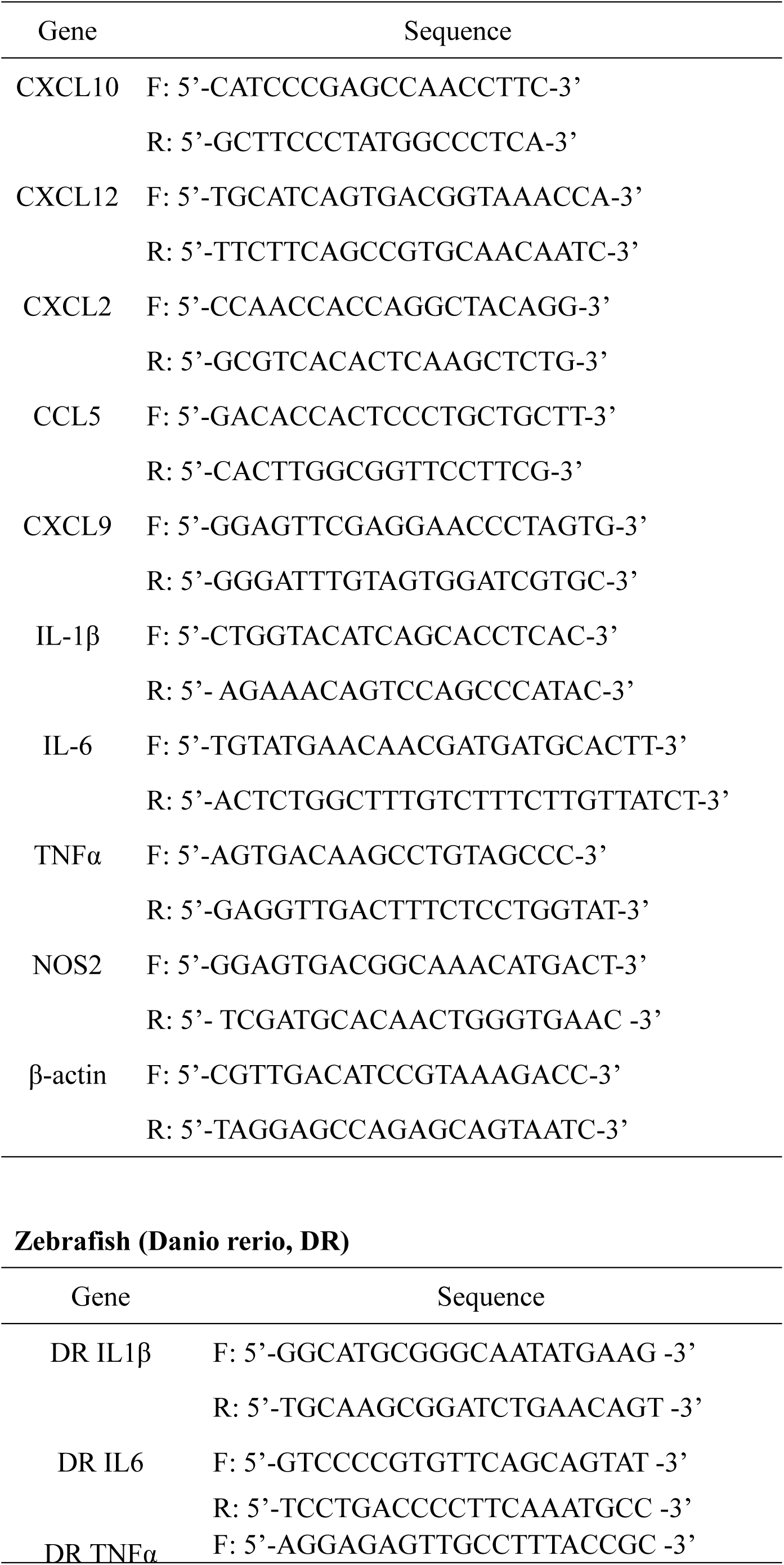
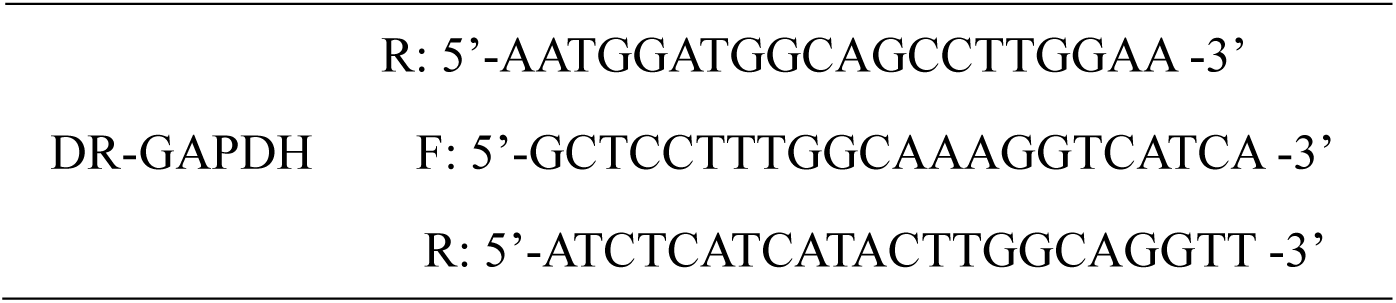
Primer pairs for real-time PCR (mouse)

RNA preparation, library construction and sequencing on BGISEQ-500 were performed at the Beijing Genomics Institute (BGI). Gene expression levels were quantified by the software package RSEM. NOISeq method was used to screen differentially expressed genes. Heat maps were generated using Excel. GO and KEGG databases were used to extrapolate differentially expressed pathways. The RNA-seq data in this study were available upon requested.

### Plasmids and siRNA transfection

Human JAK1, JAK2, JAK3, and TYK2 cDNA was obtained from 293T cell line (the sequence listed in Table 2) and was inserted into the mammalian expression vector Migr1, containing three HA tag sequence at the N-terminal region. Site-directed mutagenesis (L932A/R980A/N981A) was performed with Fast MultiSite Mutagenesis System (TransGen Biotech, cat: # FM201-01) using 3×His tag-JAK2 as a template. Plasmids were into transfected into 293T by using Lipofectamine 2000 reagent (Invitrogen, CAT: # 11668-027) ^29^. siRNAs were transfected into PEMs by using Lipofectamine® RNAiMAX Transfection Reagen (cat. # 13778075). Two JAK2 siRNA sequences were listed in Table 3.

**Table 2.**
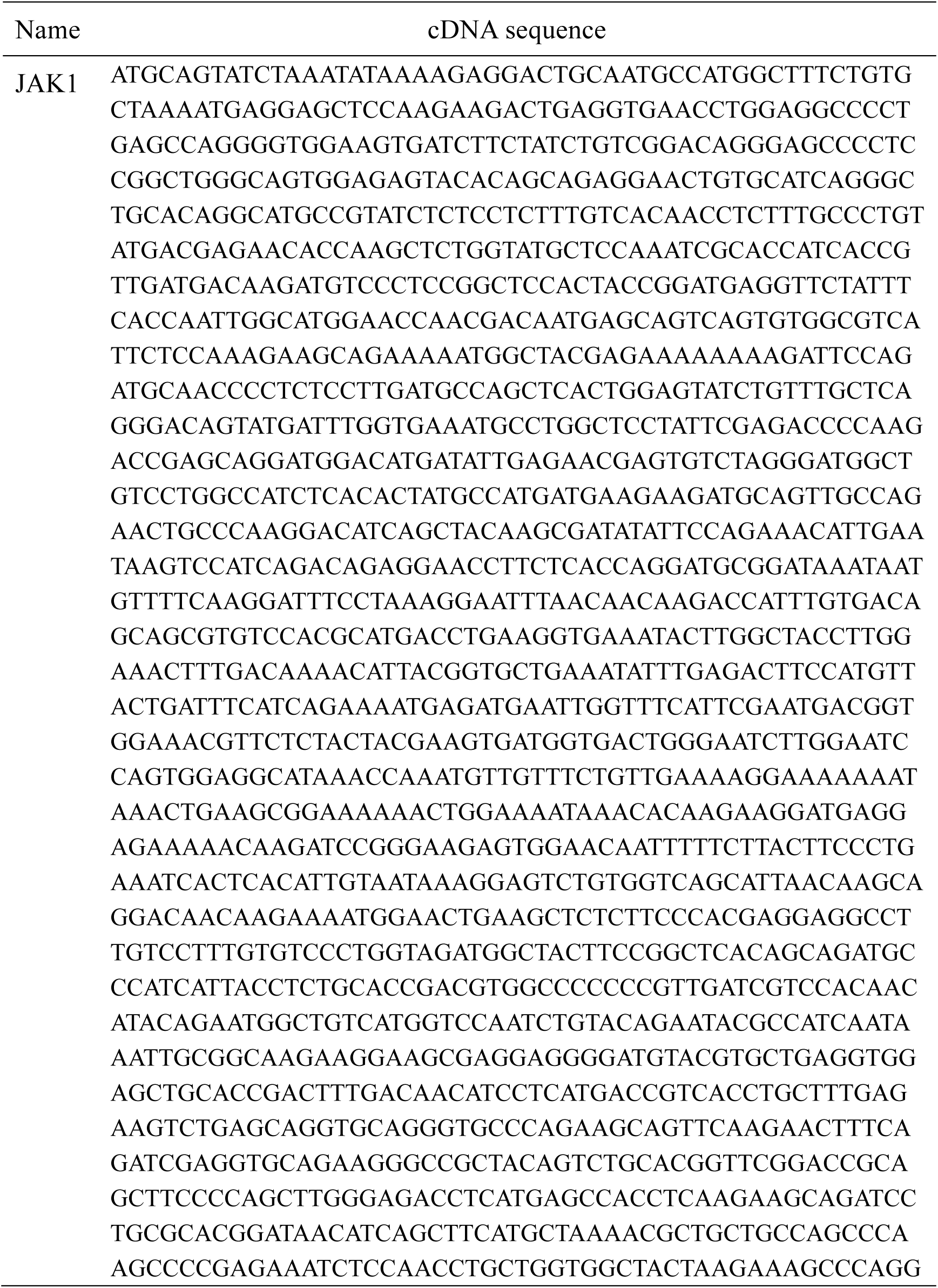
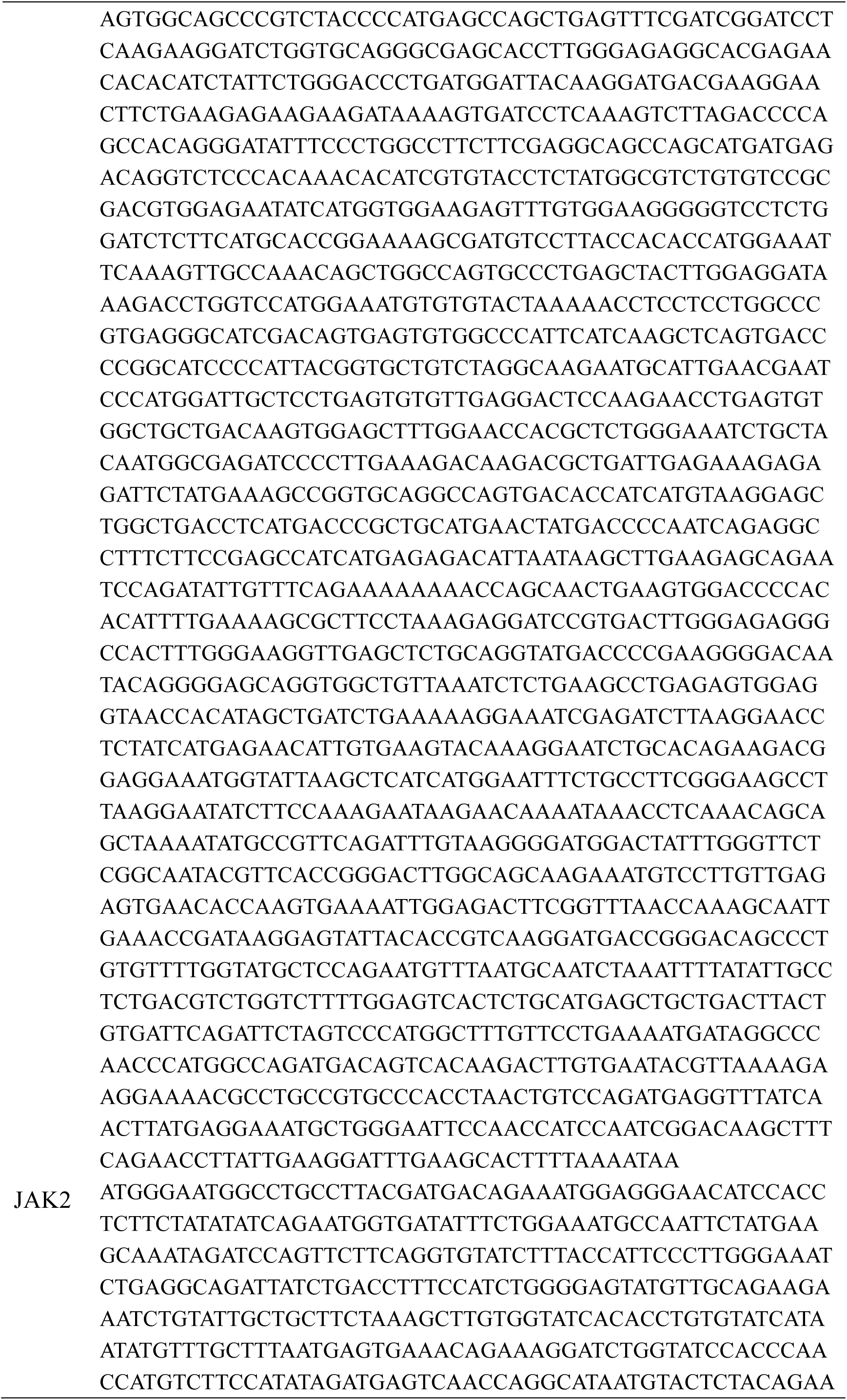
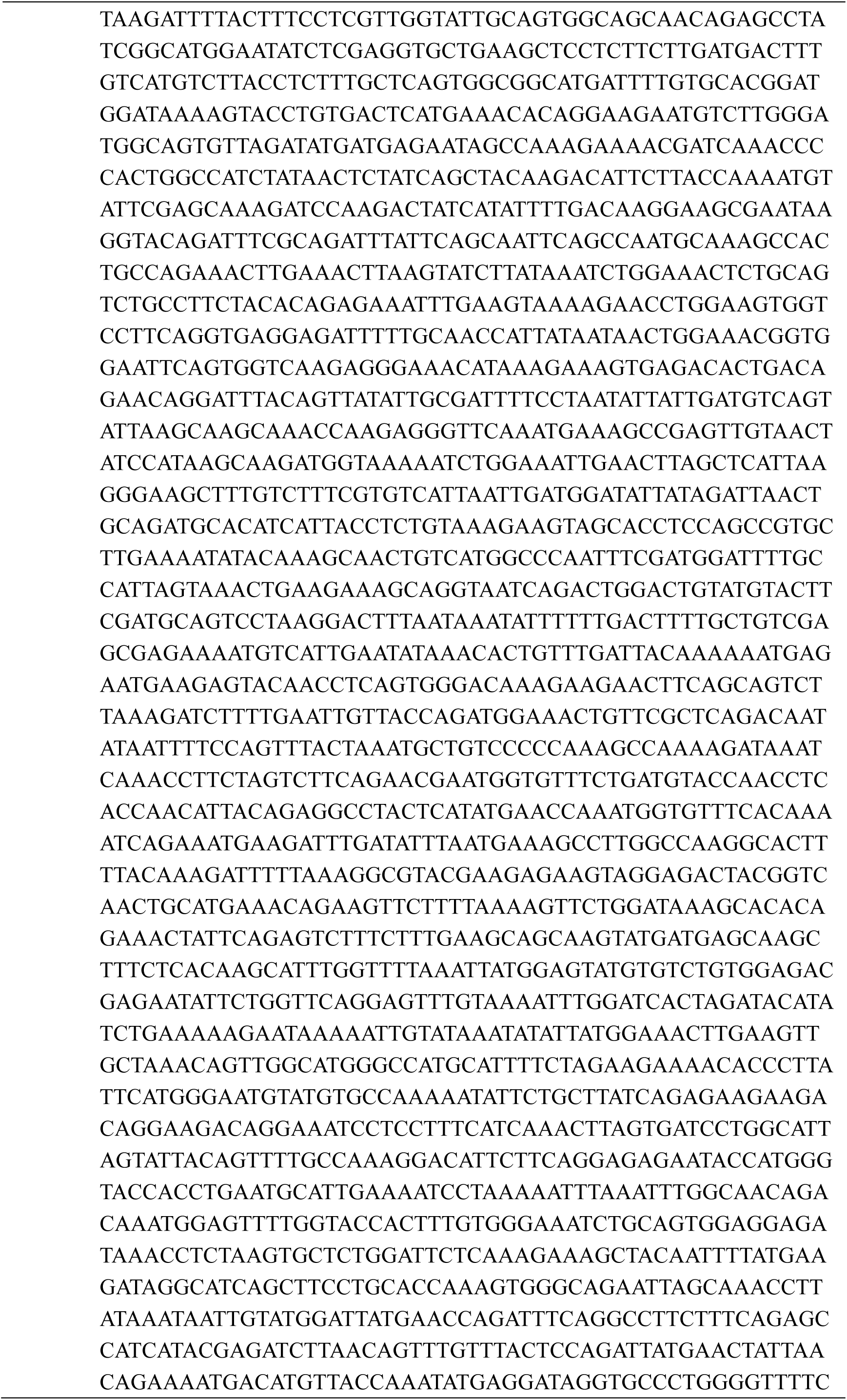
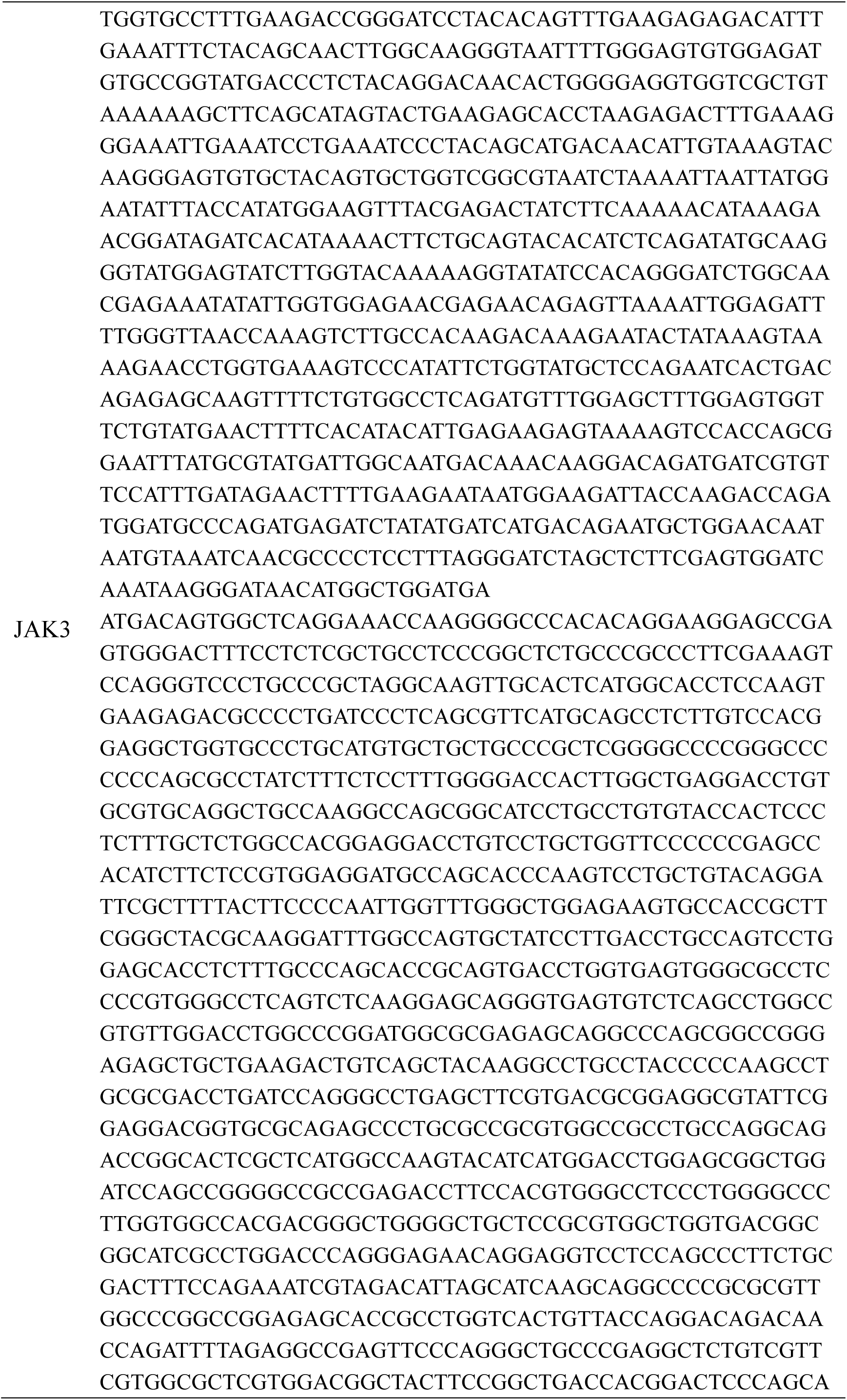
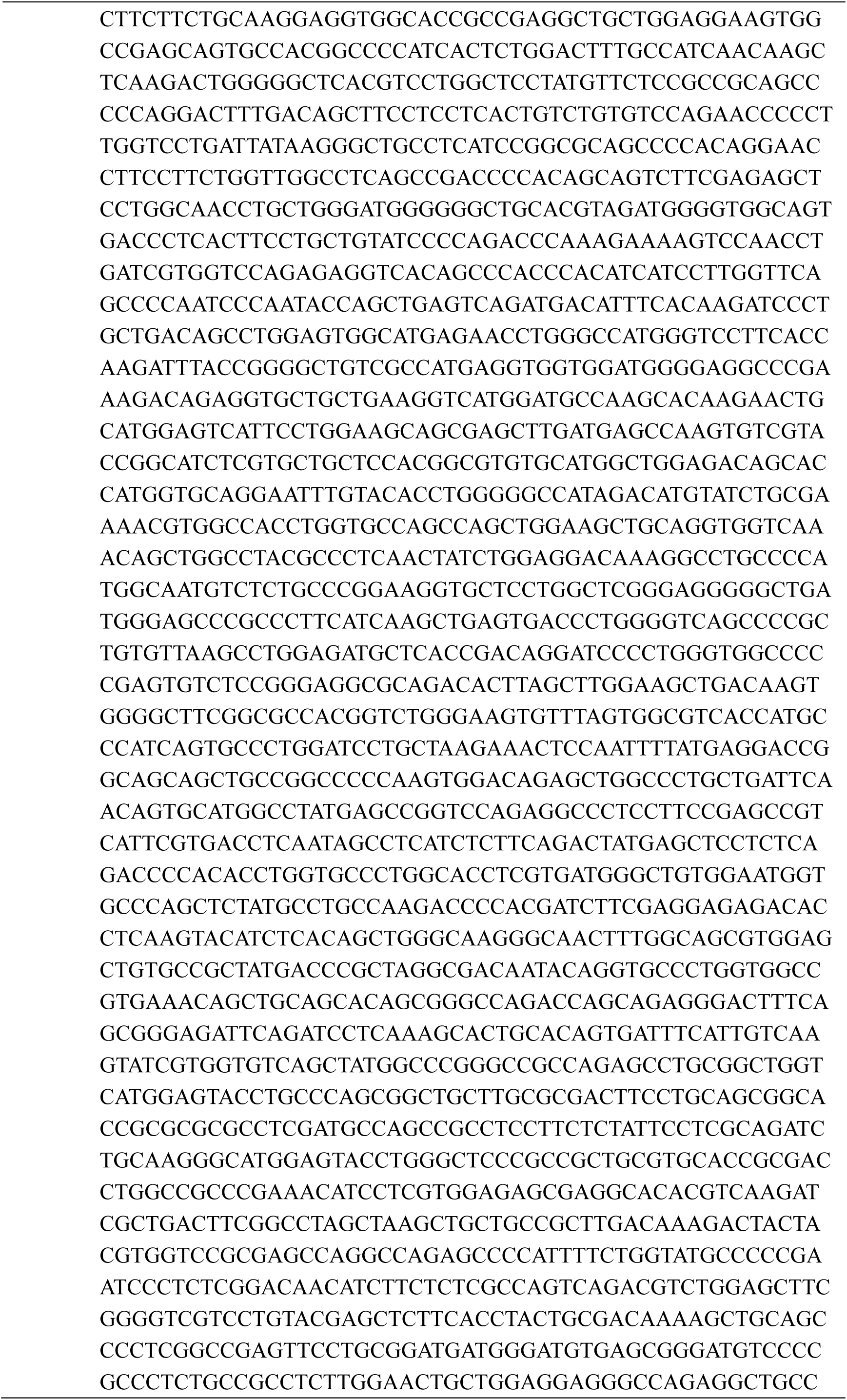
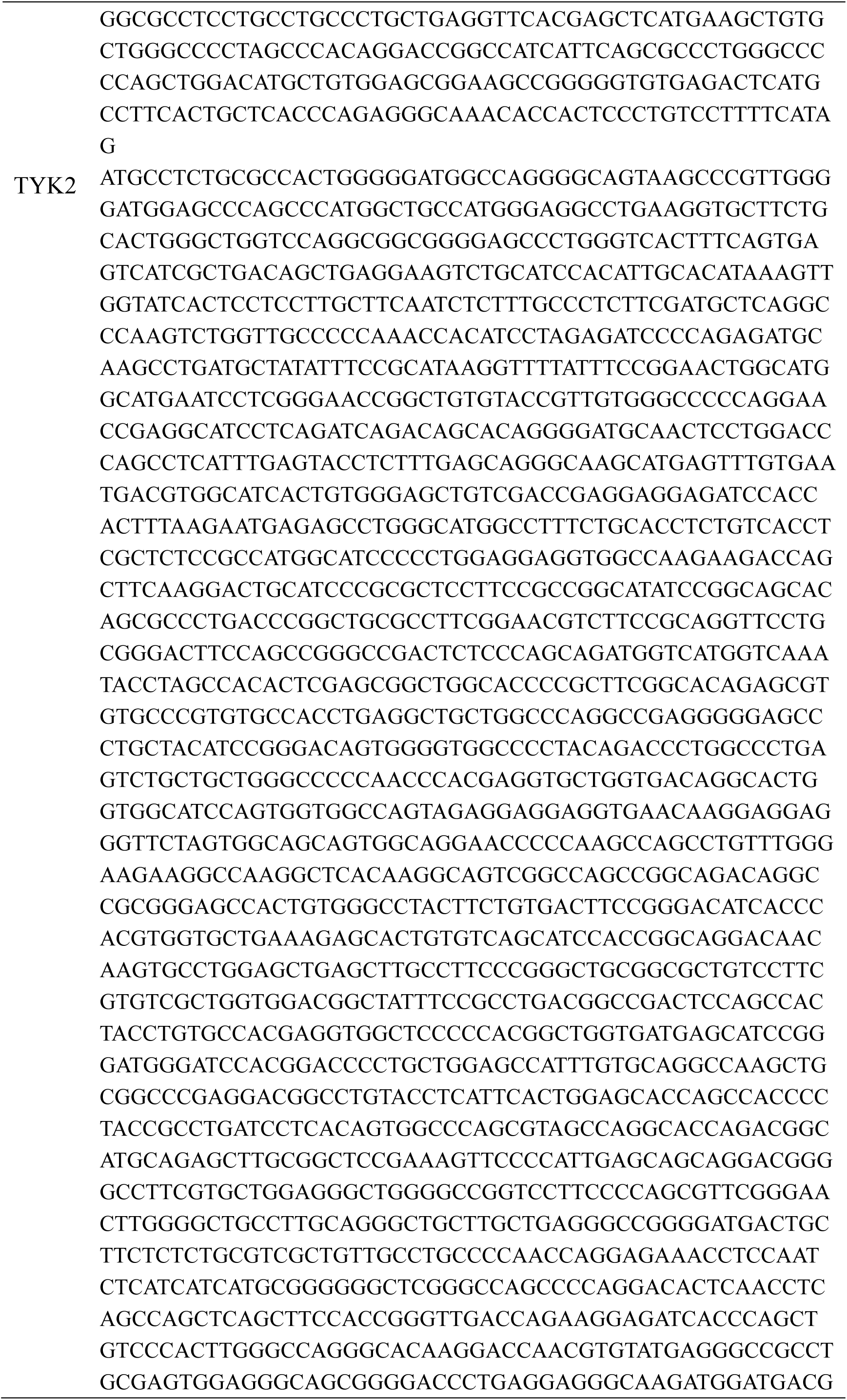
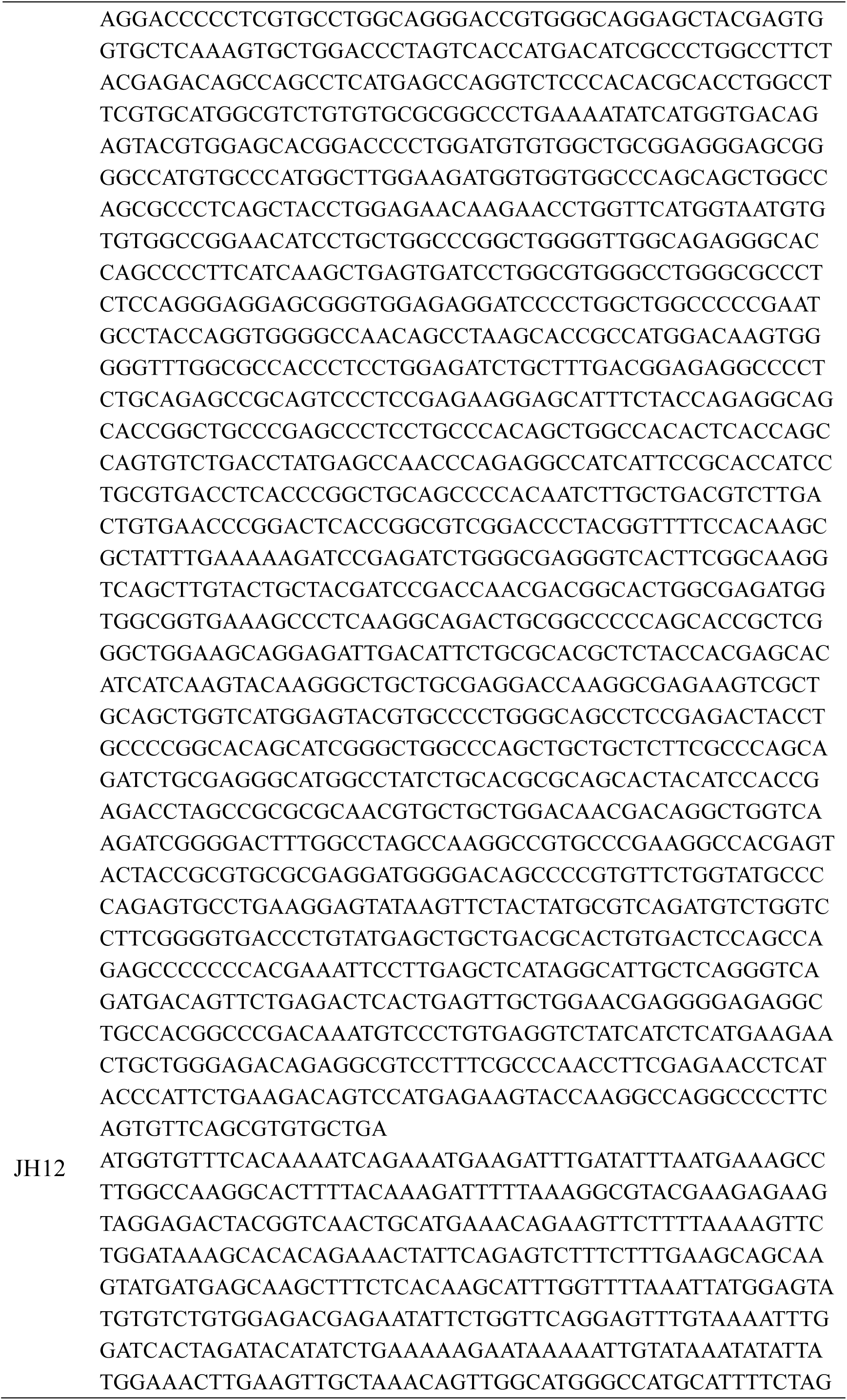
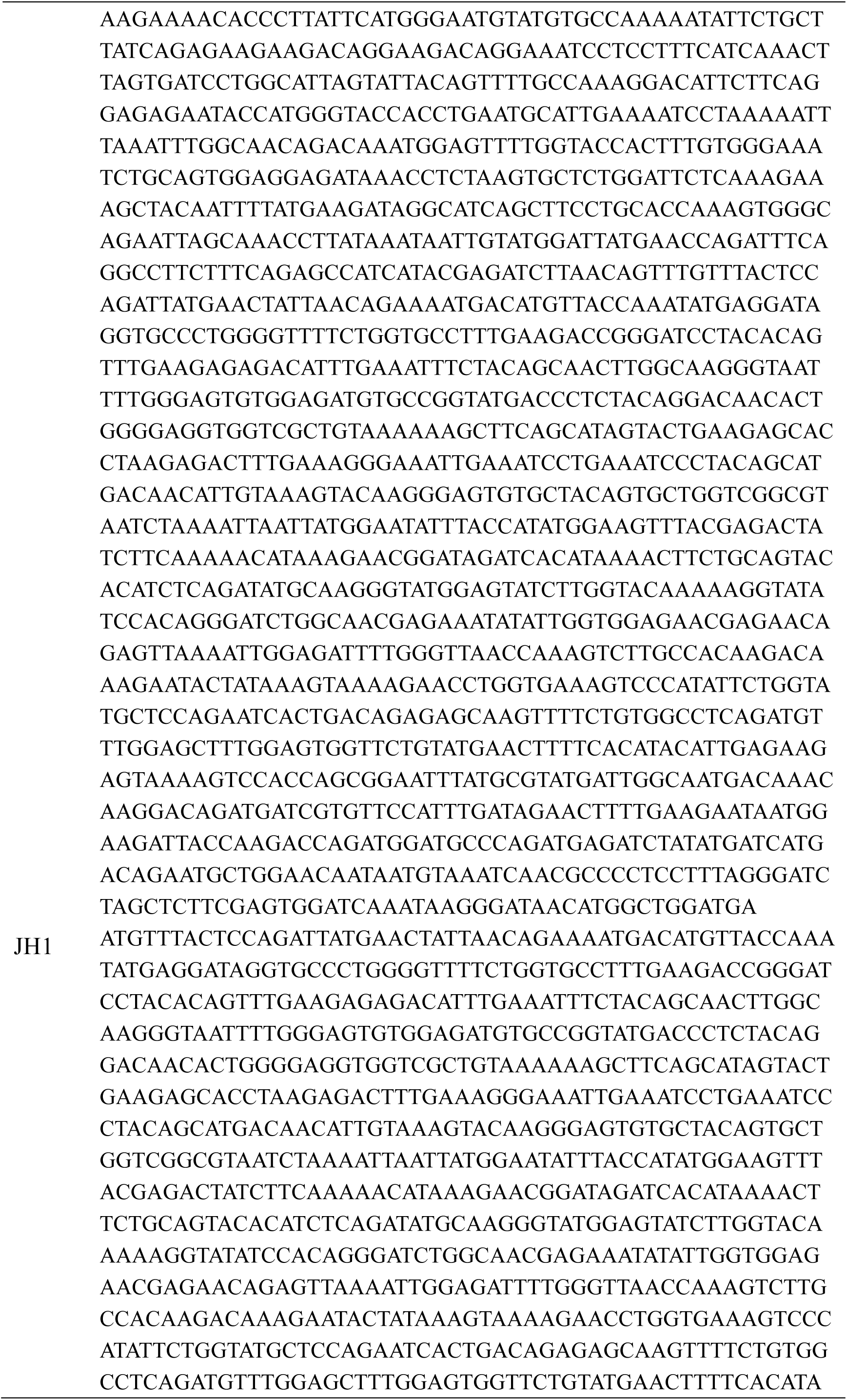
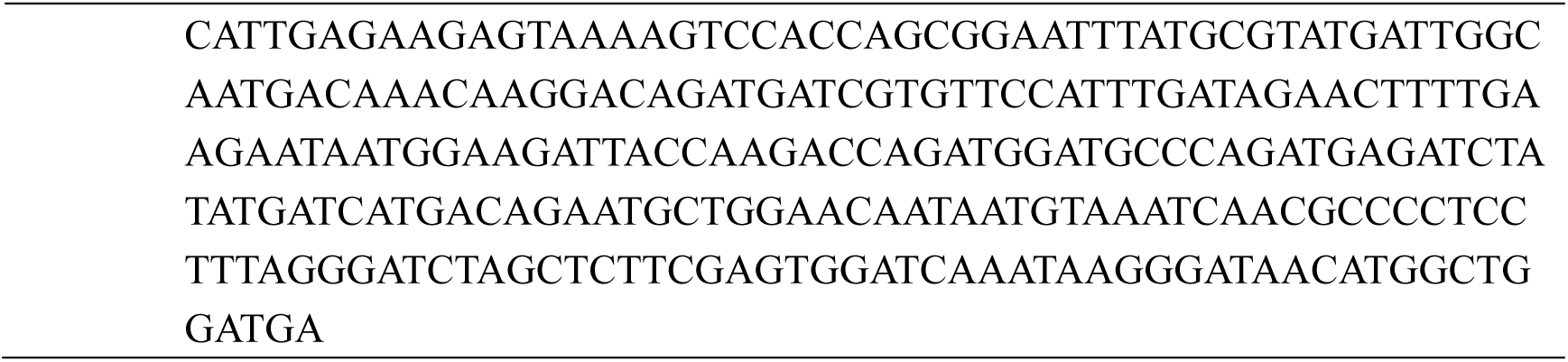
Human JAK1/2/3, TYK2, JH12 and JH1 cDNA sequence

**Table 3.**
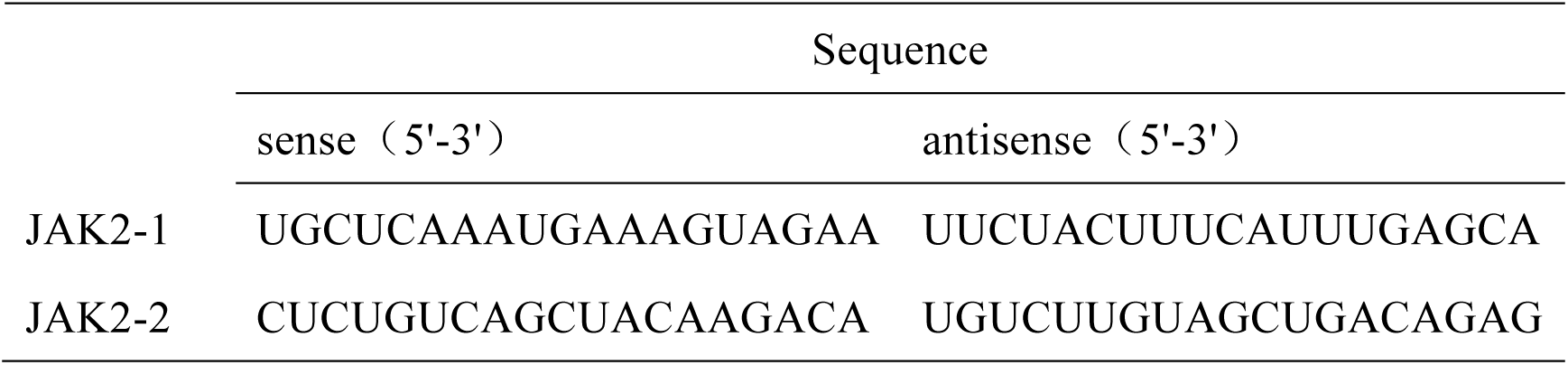
siRNA sequences for transfection

### Luciferase Assay

STAT3 and STAT5 luciferase reporter plasmids (pGMSTAT3-Lu and pGMSTAT5-Lu) were purchased from Shanghai Yeasen Biological Technology Co., Ltd. (cat: # 11503ES03 and 11554ES03). pRL-SV40-C (Beyotime Biotechnology, cat: # D2768-1μg) was used as a Renilla reporter plasmid. Migr1-3×HA-JAK1/2/3, Migr1-3×HA-TYK2 or the empty vector was transfected into 293T cells together with Renilla, pGM-STAT3-Lu or pGM-STAT5-Lu to measure the relative luciferase by the Dual-Luciferase® Reporter Assay System (Promega, CAT: # E1910).

### Protein purification of His-tagged JH1 or JH1-3A

The amino acids sequence from 808 to 1129 of the JH1 domain in human JAK2 and its cDNA sequence were shown in Table 3. JH1 or JH1-3A (L932A/R980A/N981A) was cloned into the mammalian expression vector pET-28a-c (a gift from Dr. Ronggui, Hu, Shanghai Institute of Biochemistry and Cell Biology), which contained 6 ×His tag at the N-terminal region, followed by transfection into Transetta (DE3) Chemically Competent Cell (TransGen Biotech, cat: # CD801). The Competent Cells were induced by 250 μM Isopropyl-β-D-thiogalactopyranoside (IPTG) at 16 degree. After denaturation and renaturation by gradient urea, his-tag JH1 and JH1-3A were purified by BeyoGold™ His-tag Purification Resin (Beyotime Biotechnology, cat:# P2210), which were further purified by AKATA using anion exchange column (Thermo Scientific™ Dionex™ DNAPac™ PA100, cat:# 088765).

### FACS staining and Western blot analysis

BMDMs or PEMs were suspended into 50μL Annexin V binding buffer and 1μL Anti-Annexin V-APC (eBioscience, cat: # BMS306APC-20) and incubated for 40 mins at 4℃. The cells were washed with 500 μL FACS (PBS, 0.1% sodium azide, 2% FBS) buffer and re-suspended with 50 μL FACS buffer and 1μL propidium iodide (PI, 1 mg/ml, Sigma, cat: # P4170), followed by addition of 200 μL FACS buffer into each tube. Samples were tested by Accuri C6 (BD Biosciences, San Jose, CA, USA). For WB analysis, cells were lysed with RIPA buffer (50 mM Tris-HCl pH 7.5, 150 mM NaCl, 1 mM EDTA, 1% Triton X-100, 0.1% SDS and 10% glycerol) supplemented with protease inhibitor cocktail (Roche) and 1mM PMSF, 250mM sodium fluoride, 50mM sodium orthovanadate. After centrifugation, lysates were separated by SDS-PAGE on 10% polyacrylamide gels and transferred onto Polyvinylidene fluoride (PVDF) membrane. After blocking in Tris-buffered saline and Tween 20 (TBST) with 5% BSA, the membrane was incubated with primary antibodies, washed, and then incubated with horseradish peroxidase-labeled secondary antibodies. Chemiluminescent signals were detected by MiniChemi from SAGECREATION, Beijing, China by ECL substrate (Thermo Scientific, West Palm Beach, FL, USA). The antibodies were listed in Table 4.

**Table 4.** Primary and secondary antibodies

| Name | Vendors | Number |
| --- | --- | --- |
| Anti-F4/80 antibody [CI:A3-1] | Abcam | ab6640 |
| Anti-iNOS antibody | Abcam | ab15323 |
| Phospho-IKKα (Ser176)/IKKβ (Ser177) Rabbit mAb | CST | 2688 |
| Phospho-NF-κB p65 (Ser536) (93H1) Rabbit mAb | CST | 3033 |
| Phospho-p44/42 MAPK (Erk1/2) (Thr202/Tyr204) (D13.14.4E) XP$\text{\textcircled{R}}$ Rabbit mAb | CST | 4370 |
| Phospho-p38 MAPK (Thr180/Tyr182) (D3F9) XP$\text{\textcircled{R}}$ Rabbit mAb | CST | 4511 |
| β-actin (8H10D10) Mouse mAb (HRP Conjugate) | CST | 12262 |
| IκBα Antibody | CST | 9242 |
| Phospho-Jak1 (Tyr1034/1035) Antibody | CST | 3331 |
| Phospho-Jak2 (Tyr1007/1008) Antibody | CST | 3771 |
| Phospho-Stat3 (Tyr705) (D3A7) XP$\text{\textcircled{R}}$ Rabbit mAb | CST | 9145 |
| Phospho-Stat5 (Tyr694) (D47E7) XP$\text{\textcircled{R}}$ Rabbit mAb | CST | 4322 |
| GAPDH (D16H11) XP$\text{\textcircled{R}}$ Rabbit mAb | CST | 5174 |
| HA-Tag (C29F4) Rabbit mAb | CST | 3724 |
| α-Tubulin (11H10) Rabbit mAb | CST | 2125 |
| Jak2 (D2E12) XP® Rabbit mAb | CST | 3230 |
| Anti-mouse IgG, HRP-linked Antibody | CST | 7076 |
| Anti-rabbit IgG, HRP-linked Antibody | CST | 7074 |
| Goat anti-rabbit IgG H&L (Alexa Fluor® 488) | Abcam | (ab150077) |
| Goat Anti-Mouse IgG H&L (Cy3 ®) preadsorbed | Abcam | ab97035 |

### Statistical analyses

Statistical analysis was conducted by Graphpad Prism (Version 6.0). Data represented as mean ± standard error. Statistical significance was determined by unpaired two-tailed Student’s *t-*test (or nonparametric test) with 95% confidence intervals for two groups comparison or one-way Analysis of variance (ANOVA) or nonparametric for 3 groups. As to more than 2 groups with different times points, we applied two-way ANOVA (or nonparametric) comparison. Statistical differences were indicated as \**P* < 0.05, \*\**P* < 0.01 and \*\*\**P* < 0.001.

## Results

### Intraperitoneal or oral administration of NOT protects both DBA/1J and C57/BL6 mice from collagen-induced arthritis

To explore the role of NOT in inflammatory arthritis, we generated a twice immunization collagen induced arthritis (CIA) model in DBA/1J mice (day 0 and day 21), followed by intraperitoneal injection with NOT or the saline control at day 22 after the first immunization (Fig. 1a). We observed that NOT treatment significantly reduced the clinical scores of the CIA mice, but showed similar weight as those CIA mice treated with the saline control (Fig. 1b). Three-dimensional micro-CT graphic examination showed that the saline-treated CIA mice reduced the Talus bone volume (blue) with severer bone erosion (i.e. bone destruction), while the NOT-treated CIA mice recovered the phenotype of Talus bone volume and also reduced bone erosion to a similar level as the mock mice (Fig. 1c). ABOG staining data have revealed the increased Synovitis (including Talus and Meniscus synovitis) and reduced cartilage (including Talus and Meniscus cartilage) area in the saline-treated CIA mice. By contrast, NOT treatment significantly reduced synovitis and increased cartilage area (Fig. 1d). In addition, we observed that increasing number of osteoclast in Taus or Meniscus in the saline-treated CIA mice, while NOT treatment could markedly reduce the numbers of osteoclast in Taus or Meniscus when compare to the saline group via Trap staining (Fig. 1e).

**Fig. 1.**
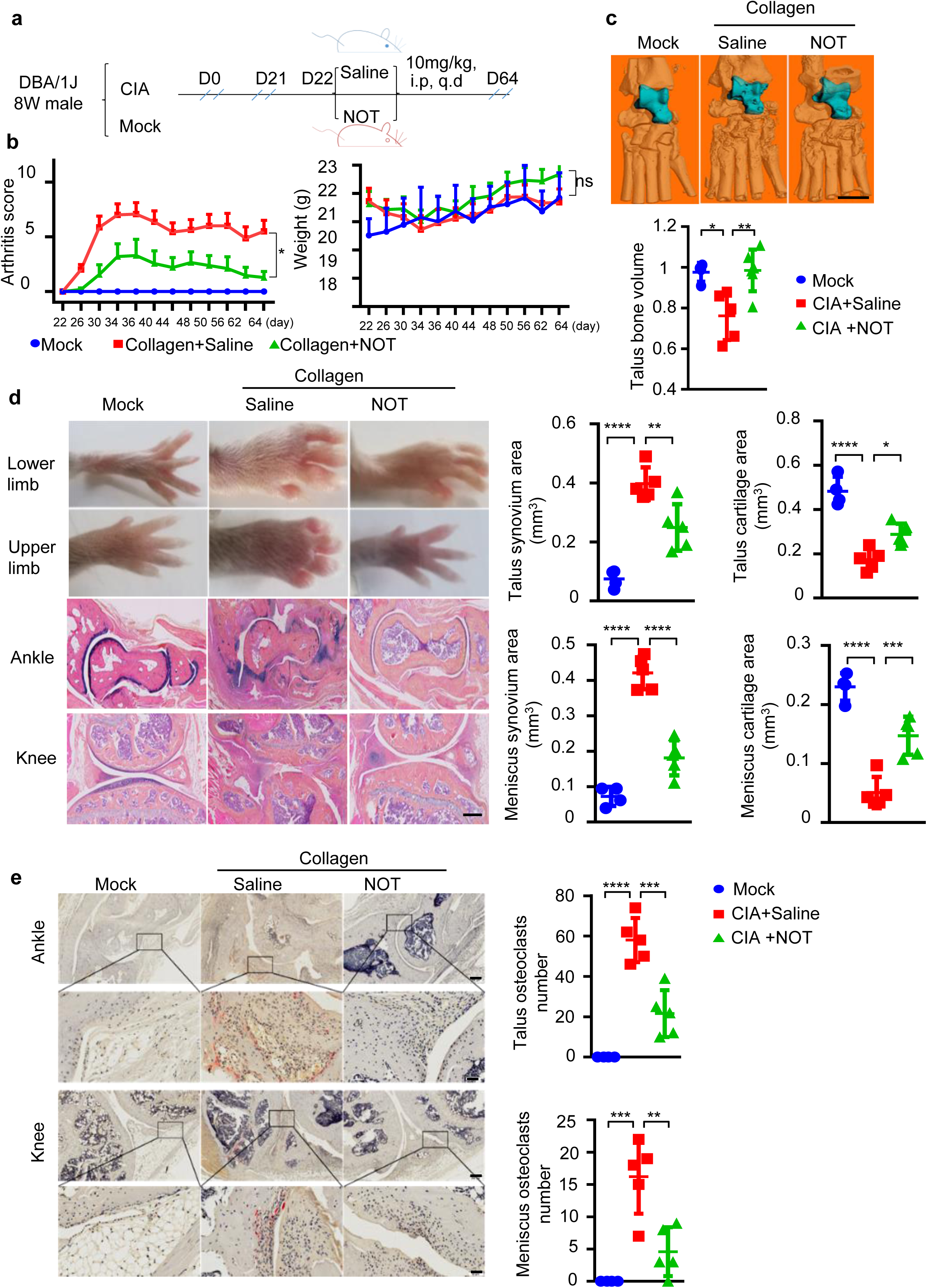
Intraperitoneal injection of NOT protects DBA/IJ mice from collagen-induced arthritis. **a-b**, A schematic diagram illustrating the procedure of NOT treatment or the saline vehicle treatment in male DBA/1J mice that were i.p. injected with collagen to induce RA (i.p. intraperitoneal; q.d. once a day). Clinical arthritic scores (left) and body weight (right) in the non-CIA mock (n=4), the CIA mice treated with NOT (n=14) or the CIA mice treated with saline (n=11). Data represent mean ± SEM from two experiments. Two-way ANOVA. **c**, Representative μCT images of ankle joints (upper panel, Scale bar: 1 mm) and quantification of talus bone volume (lower panel). Data represent mean ± SEM from two experiments (n=4 Mock, n=5 Saline, n=6 NOT). One-way ANOVA. **d**, Representative images of lower and upper limbs, and representative photomicrographs of ABOG-stained tissue sections of ankle and knee joints (Scale bar: 100 μm). Quantification of histological analysis of synovitis and cartilage area around talus and meniscus of each group (n=4 Mock, n=5 Saline, n=5 NOT). Data represent mean ± SEM from two experiments, one-way ANOVA. **e**, Representative images of TRAP staining to show osteoclasts from synovial tissues around talus and meniscus in the hind paw joints (left; Upper scale bar: 200 μm, Lower bar: 50 μm). The numbers of osteoclasts in synovial tissues around talus and meniscus (right, n=4 Mock, n=5 Saline, n=5 NOT). Data represent mean ± SEM, one way ANOVA. *p<0.05, **p<0.01, ***p<0.001, ****p<0.0001.

To access the effect of NOT via oral administration, we next generated the CIA model by one immunization in DBA/1J mice followed by the treatment with NOT that was dissolved in CMC-Na or the CMC-Na control at day 42 (i.e. the onset of symptoms) via a route of oral gavage (Supplementary Fig. 1a). Similar to the effect of intraperitoneal (i.p) injection, NOT treatment significantly improved the clinical symptoms, including reducing the ameliorated arthritis score without affecting the body weight (Supplementary Fig. 1b), decreasing synovitis and increased cartilage area (including Talus and Meniscus sites) (Supplementary Fig. 1c) and decreasing synovial inflammation and less structural damages of the cartilage and bone (Supplementary Fig. 1d).

To further verify the effect of NOT using other mouse strains, we generated the twice immunization CIA model in C57/BL6 mice (day 0 and day 21), followed by i.p injection of NOT or the saline control (Fig. 2a). We observed that NOT treatment could also showing similar therapeutic effect as those observed in the DBA/1J model and protect C57/BL6 mice from the CIA-induced RA (Fig. 1). The NOT treatment significantly reduced the clinical scores (Fig. 2b), decreasing synovitis and increased cartilage area (including Talus and Meniscus sites) (Fig. 2c), lessened the destruction of cartilage and bone (Fig. 2d) and reducing the numbers of osteoclasts from Talus and Meniscus (Fig. 2e).

**Fig. 2.**
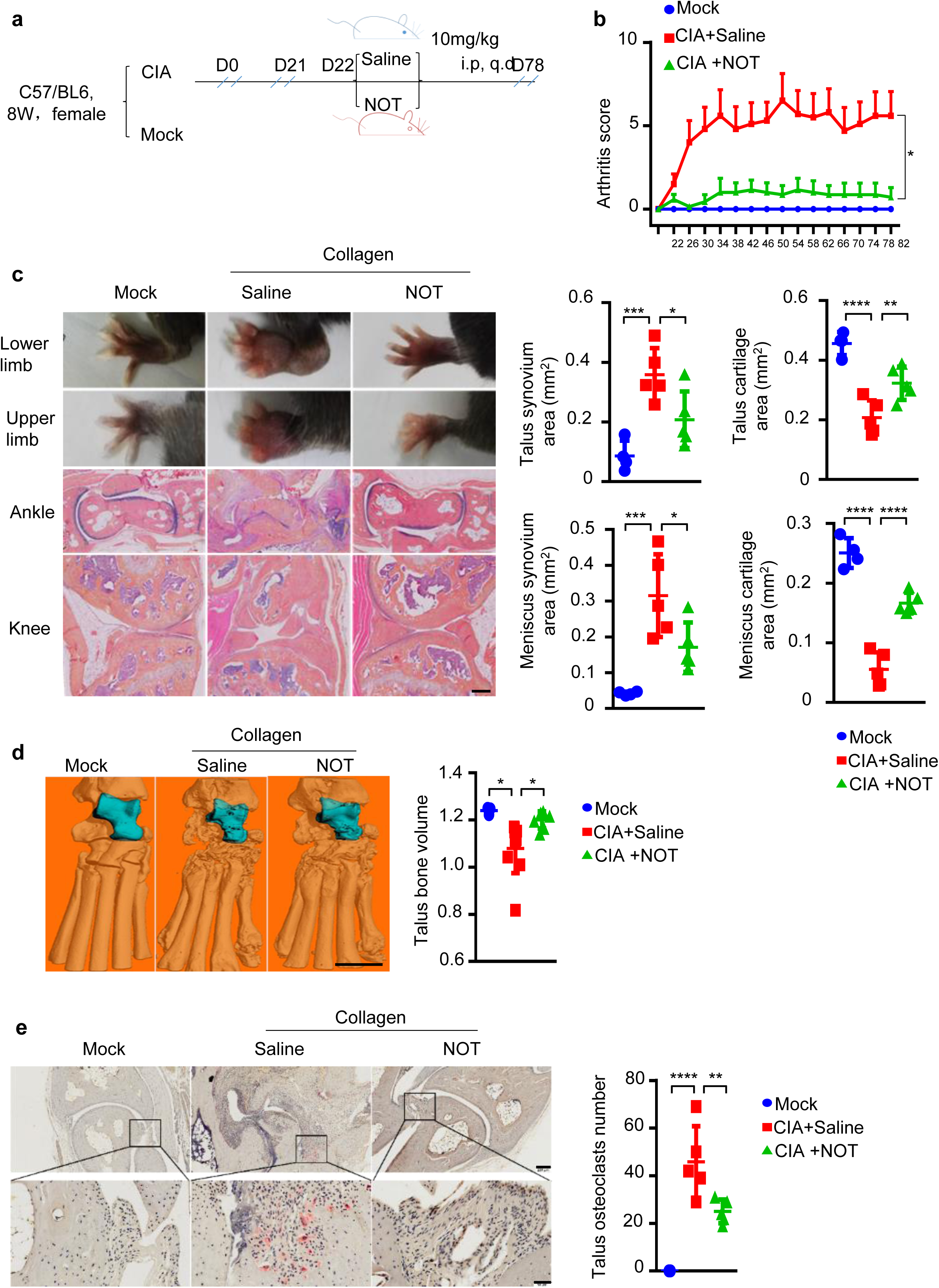
Intraperitoneal injection of NOT protects C57/BL6 mice from collagen-induced arthritis. **a-b**, C57BL/6 mice were immunized twice with collagen to induce CIA and the treatment with NOT (n=10) or Saline (n=7) were performed as indicated. The unimmunized mock mice were used as control (n=4). The clinical score was measured. Data represent mean ± SEM, two-way ANOVA. **c**, Representative photographs of lower limbs and upper limbs and representative images of ABOG-stained tissue sections of ankle or knee joints. Quantification of histological analysis of synovitis and cartilage area around talus and meniscus of each group (Scale bar:100 μm; n=4 Mock, n=5 Saline, n=5 NOT). Data represent mean ± SD, one-way ANOVA. **d**, Representative microCT images (Scale bar: 1mm) and quantification of talus bone volume were calculated (n=3 Mock, n=10 Saline, n=7 NOT). Data represent mean ± SD, one-way ANOVA. **e**, Representative images of TRAP staining to show osteoclasts from synovial tissues around talus and meniscus (Upper scale bar: 200 μm, Lower bar: 50 μm). The numbers of osteoclasts was analyzed (n=4 Mock, n=5 Saline, n=5 NOT). Data represent mean ± SD, one way ANOVA. *p<0.05, **p<0.01, ***p<0.001, ****p<0.0001.

Moreover, we observed no obvious toxicity from the NOT treated mice when compared to the untreated mice, including concentrations of ALT, AST and Urea, CRE (Supplementary Fig. 2a,b). Furthermore, hematoxylin and eosin (H&E) staining showed no damages in the liver, kidney, lung, and spleen (Supplementary Fig. 2c) from the NOT-treated or untreated mice.

### NOT inhibits production of multiple pro-inflammatory cytokines and chemokines in TNFα- and LPS-treated macrophages

Previous studies and our data have demonstrated that elevated macrophages infiltration and enhanced levels of inflammatory cytokines are correlated with the severity of rheumatoid arthritis (RA) ^30, 31, 32^ (Fig. 3a-c). We next asked whether NOT treatment affected the *in vivo* macrophage infiltration and production of inflammatory cytokines. Using F4/80+/iNOS+ as the markers to identify inflammatory macrophages, we found that NOT treatment markedly decreased the numbers of F4/80+/iNOS+ macrophages in synovial of the CIA mice when compared to the Saline control-treated CIA mice (Fig. 3a). In addition, NOT treatment substantially suppressed IL-1β, TNFα and IL-6 levels in serum (Fig. 3b) as well as in synovial fluid (Fig. 3c), which was restored to similar levels as those from the non-collagen induced mock mice.

**Fig. 3.**
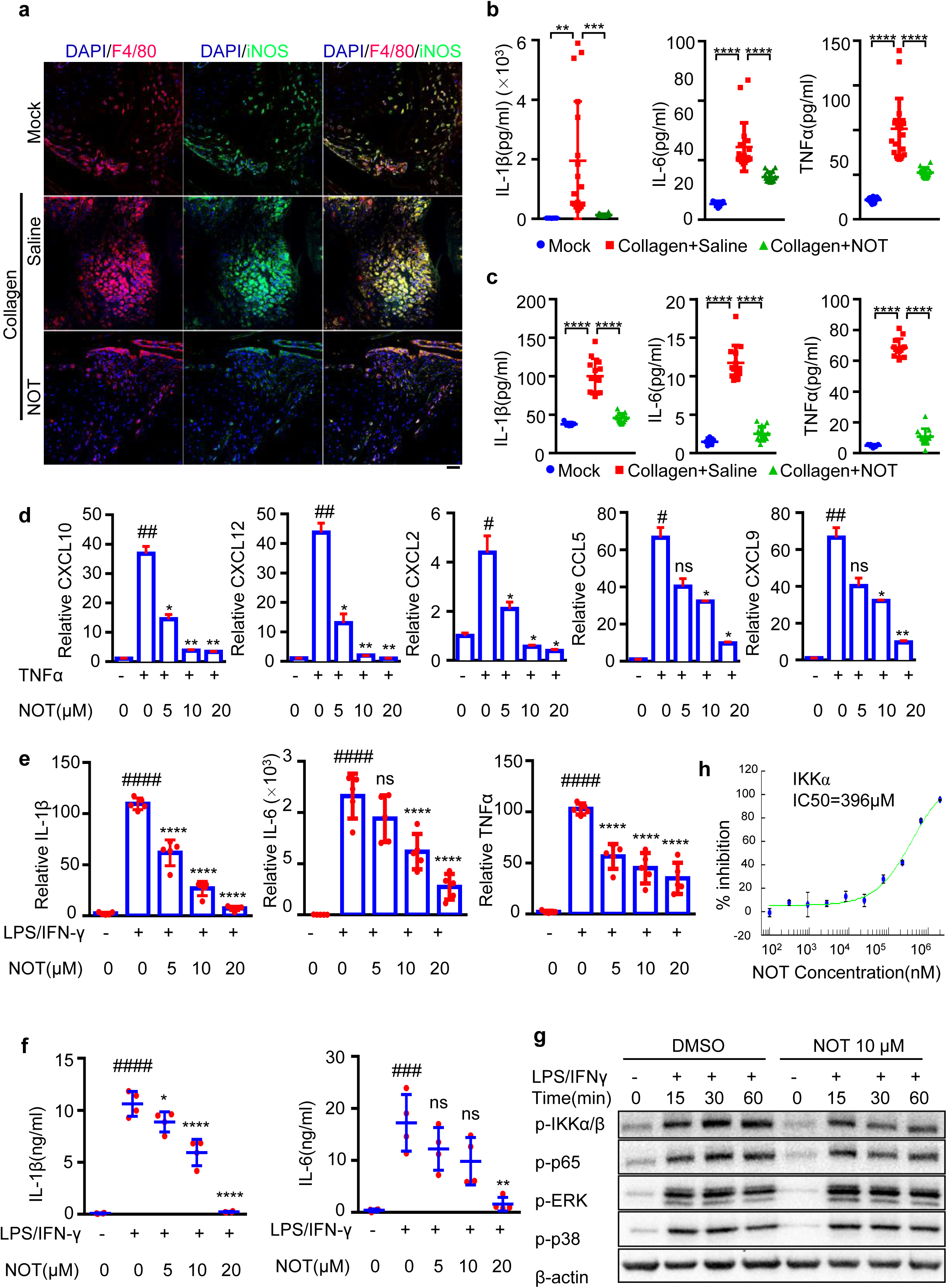
NOT inhibits production of multiple pro-inflammatory cytokines and chemokines in TNFα- and LPS/IFNγ-treated macrophages. **a**, Representative immunofluorescence images (Scale bar: 50 μm) show F4/80+ (red) and iNOS+ (green) cells in synovium around talus (n=3 Mock, n=7 Saline, n=7 NOT). **b**, Serum IL-1β, IL-6, and TNF-α concentrations of the mock (n=9) and the CIA mice treated with NOT (n=18) or saline (n=18). Data represents mean ± SD of three experiments, one-way ANOVA, **p<0.01, ***p<0.001, ****p<0.0001. **c**, IL-1β, IL-6, and TNF-α concentrations in the posterior limb (about 0.5 cm from the tibiotalar joint to the tip toe) were measured from the mock (n=5) or the CIA mice treated with NOT (n=13) or saline (n=13). Data represents mean ± SEM of three experiments, one-way ANOVA. ****p<0.0001. **d-e**, The mRNA levels of CXCL10, CXCL12, CXCL2, CCL5 and CXCL9 were checked in TNFα (100 ng/ml, 6 hrs)-treated **(d)** or LPS (1 µg/ml)/IFN-γ (100 ng/ml)-treated **(e)** BMDMs in the absence or presence of NOT (0, 5, or 10 or 20 µM, 3 hrs). Data represents mean ± SD of at least 3 experiments, one-way ANOVA. **f**, IL-6 concentrations were measured with ELISA in the supernatants of LPS/IFN-γ-treated BMDMs in the absence or presence of NOT (0, 5, or 10 or 20 µM, 3 hrs). To measure IL-1β concentrations, BMDMs were treated with LPS for 6 hrs followed with SL1344 infection for 0.5 hrs in the absence or presence of NOT. Data represents mean ± SD of four experiments. One-way ANOVA. #P<0.05, ##P<0.01, ###P<0.001,####P<0.0001 were labeled when TNFα- or LPS/IFNγ-treated samples were compared with the resting Mock group (no TNFα or LPS/IFN-γ treatment). And ns indicating P>0.05, *P<0.05, **P<0.01, ****P<0.0001 were labeled when the NOT-treated samples were compared with TNFα- or LPS/IFNγ-treated samples in **d-f**. **g**, The phosphorylation levels of IKKα/β, p65, ERK, p38 and the protein level of β-actin were checked by immunoblotting in cell lysates of BMDMs after stimulated with LPS(1µg/ml)/IFN-γ (100 ng/ml) for 15, 30, and 60 min in the absence or presence of NOT (10µM, pretreated for 3hrs). Representative blots of three independent experiments. **h**, The effect of NOT to inhibit IKKα activation was measured by Z’-Lyte assay.

To determine whether NOT directly targets macrophages to inhibit the production of inflammatory cytokines, we used TNFα or LPS/IFNγ to stimulate macrophages *in vitro* with or without NOT treatment. Notably, NOT showed a dose-dependent effect to reduce the mRNA levels of CXCL10, CXCL12, CXCL2, CCL5 and CXCL9 in TNFα-stimulated bone marrow-derived macrophages (BMDMs) (Fig. 3d). Moreover, NOT significantly reduced the mRNA and protein levels of IL-1β, IL-6 and TNFα in LPS/IFNγ-stimulated BMDMs (Fig. 3e,f). Importantly, treatment with NOT at the highest doses (20 μM) for 72 hours did not induce PEMs or BMDMs apoptosis and have no obvious effect on cell viability (Supplementary Fig. 3a,b).

NF-κB is one of the main transcription factors to induce expression of inflammatory cytokines and is activated by both TNFα and LPS/IFNγtreatment. In addition, NF-κB was previously reported to be involved in the development of RA^33, 34^. We found that NOT treatment reduced the phosphorylation levels of IKKα/β and p65, without any effect on the levels of pERK and p-p38 in LPS/IFNγ stimulated macrophages (Fig. 3g). To further explore how NOT treatment could reduce IKKα/β phosphorylation, we used a cell-free analysis (Z’-LYTE Kinase Assay, SelectScreen Biochemical Kinase Profiling Service from Thermo Fisher) to measure the IC50 of NOT on the NF-κB pathway related kinases via the incubation of NOT with IKKα, IKKβ, and TAK1 respectively, followed by checking the activity of IKKα, IKKβ, and TAK1. Although NOT displayed a minor inhibitory effect on the activation of IKKα, with a half maximal inhibitory concentration (IC50) reaching 396μM, this concentration was far greater than the concentration we used to treat macrophages (Fig. 3h). In addition, NOT, even when used at 500 μM, showed no inhibition on the activation of IKKβ and TAK1. These results indicate that NOT might not directly target TAK1 or IKKα/β to induce anti-inflammatory effect.

### NOT inhibits JAK-STAT activation to attenuate inflammation

To explore the mechanism of how NOT treatment inhibited pro-inflammatory cytokines production, we measured differences of gene expression via the whole genome RNA sequencing (RNA-seq) between the NOT-treated and untreated macrophages in response to TNFα stimulation. We identified 54 genes in the NOT-treated macrophages that were significantly down-regulated, including the expression levels of CXCL10, CCL5, CCL4, other chemokines or cytokines as well as the tyrosine kinase JAK2 (Fig. 4a). When the KEGG pathways were analyzed, these different genes were enriched in “TNF signaling pathway”, “Chemokine signaling pathway”, “TLR pathway” and “JAK-STAT pathway” (Fig. 4b).

**Fig. 4.**
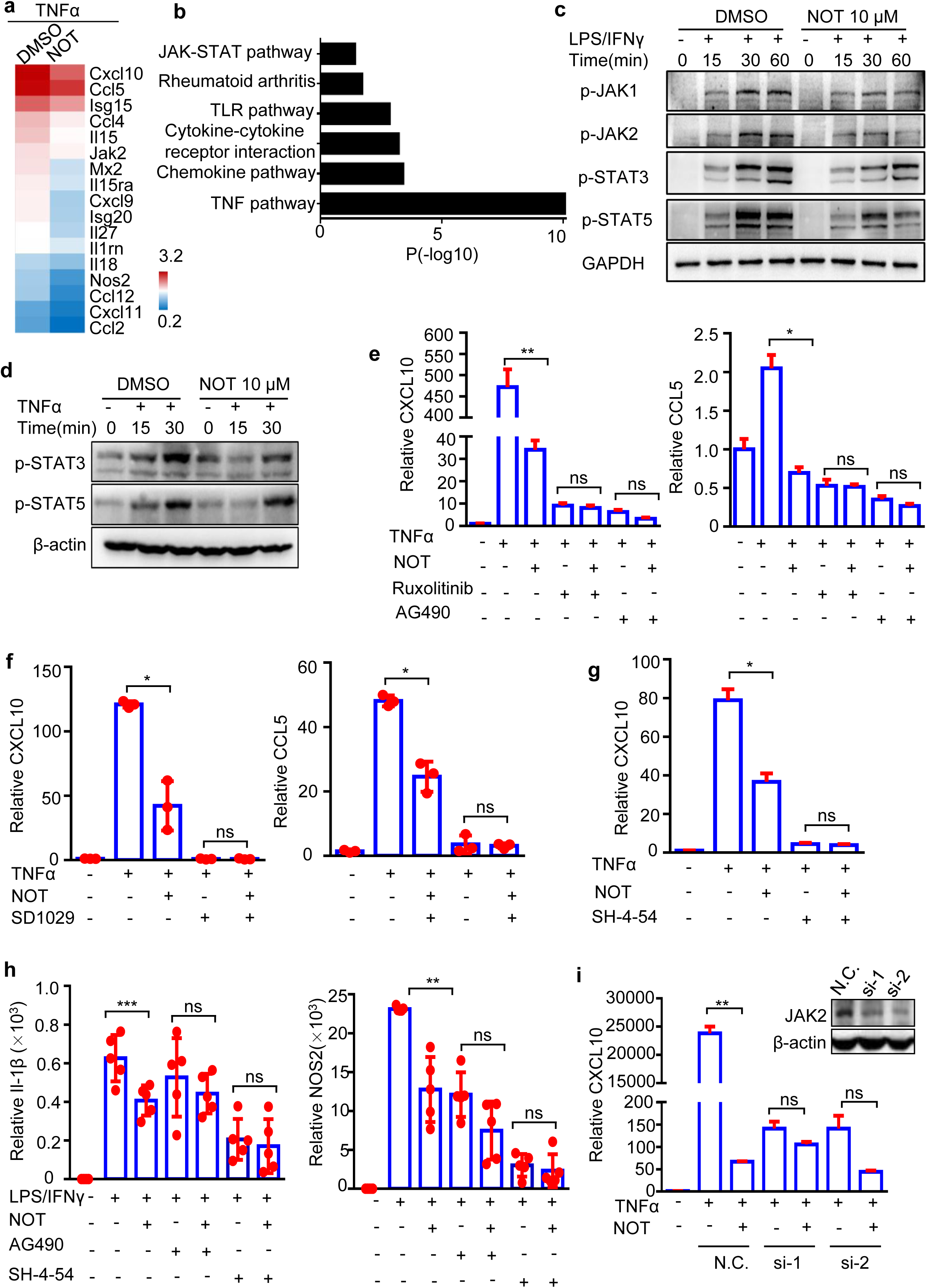
NOT inhibits JAK-STAT activation to attenuate inflammation. **a-b**, Some of the differentially expressed genes (2-folds, p≤0.05, n=3 from each group) were listed from the RNA sequence (RNA-seq) data of TNFα-stimulated BMDMs (6 hrs), which were pretreated with or without NOT (10µM, 3 hrs). Pathway analysis was determined from all differentially expressed genes (≥2-fold changes) between the DMSO and NOT treated samples by DAVID 6.8. **c-d**, The phosphorylation levels of JAK1, JAK2, STAT3, STAT5 and the protein level of GAPDH were detected by immunoblotting in cell lysates of LPS(1µg/ml)/IFN-γ (100 ng/ml)-stimulated or TNFα (100 ng/ml)-stimulated BMDMs for the indicated time points, which were pretreated with or without NOT (10µM, 3 hrs). Representative blots of three independent experiments are shown. **e-g**, The mRNA levels of CXCL10 and CCL5 were checked in TNFα (100 ng/ml)-stimulated BMDMs, which were pretreated with NOT (10µM) in the presence or absence of the JAK2 inhibitors Ruxolitinib (10µM), AG490 (50µM) **(e)**, SD1029 (10µM) **(f)** or the STAT3/5 inhibitor SH-4-54 (10µM) **(g)** for 3 hrs. Data are mean ± SEM from at least 3 experiments, one-way ANOVA. **h**, The mRNA levels of IL-1β and NOS2 were checked in LPS (1μg/ml)/IFNγ (100ng/ml)-stimulated BMDMs, which were pretreated with NOT (10µM) in the presence or absence of the JAK2 inhibitor AG490 (50µM) or the STAT3/5 inhibitor SH-4-54 (10µM) for 3 hrs. Data represents mean ± SD of five experiments, one-way ANOVA. **i**, CXCL10 mRNA levels were checked in TNFα (100 ng/ml)-stimulated PEMs, which were transfected with si-JAK2#1 and siJAK2#2 followed by pretreatment with NOT (10 µM) for 3 hrs. Representative data are mean ± SEM from at least 3 mice, one-way ANOVA. JAK2 knockdown efficiency was confirmed by immunoblotting (Upper panel). ns P>0.05, *P<0.05, **P<0.01, ****P<0.0001 were labeled when the NOT-treated samples were compared with TNFα- or LPS/IFNγ-treated samples in **e-i**.

Since the JAK-STAT pathway was identified in the NOT-treated macrophages, we next tested the phosphorylation levels of JAKs and STATs. Interestingly, NOT treatment significantly inhibited JAK1/2 phosphorylation and the downstream Stat3/5 phosphorylation induced by LPS/IFNγ or by TNFα (Fig. 4c,d). To further test whether NOT inhibited inflammation through JAK1/2 and STAT3/5, we treated macrophages with the JAK1/2 inhibitors, including SD1029, AG490 and Ruxolitinib ^35^, and the STAT3/5 inhibitor SH-4-54 ^36^. All of these inhibitors could markedly inhibit the production of CXCL10, CCL5, IL-1β, or NOS2 at mRNA or at protein levels in TNFα- or LPS/IFNγ-treated macrophages (Fig. 4, e-h). Moreover, pretreatment with these inhibitors abolished the anti-inflammation effect of NOT (Fig. 4, e-h). In addition, to further confirm the function of JAK, we knocked down (KD) JAK1/2/3 and TYK2 expression with siRNAs in macrophages.We found that the inhibitory effect of NOT was no longer displayed in JAK2/3 KD macrophages (Fig. 4i, the KD of JAK1/3 and TYK2 were not shown). Those results indicate that NOT depends on the JAK-STAT pathway to mediate the anti-inflammation effect.

### NOT inhibits JAK-STAT activity via directly binding to JAK2/3

To find the direct target proteins of NOT, We applied the Surflex-Dock analysis. The Surflex-Dock (version 2.6, http://www.tripos.com) was used to screen the potential partners for NOT against the sc-PDB (Version 2013) database that contains 9,277 protein binding sites^37, 38, 39^. Each docking pose was rescored with Gaussian shape similarity^40^ implemented in the Shape it (v1.0.1)^41^. Only the docking results with Gaussian shape Tanimoto greater than 0.68 and polar score greater than 1 were examined visually to confirm the docking pose and essential interactions. The tyrosine-protein kinase JAK2 (the gaussian similarity was 0.77 and the polar score was 2.48) was ranked as the top 7 candidate. The predicted affinity (dock score) and shape similarity between the pose and the cognate ligand were showed in Table 5. Based on this analysis, we hypothesized that NOT might interact with JAK2 to inhibit the JAK-STAT pathway.

**Table 5.** Docking result of NOT against JAK2 (PDB code:4E6Q) using Surflex-Dock POSE ID DockScore PolarScore ShapeTanimoto TverskyRef TverSkyDB

| POSE ID | DockScore | PolarScore | ShapeTanimoto | TverskyRef | TverSkyDB |
| --- | --- | --- | --- | --- | --- |
| 0 | 7.83 | 1.36 | 0.61 | 0.80 | 0.72 |
| 1 | 7.67 | 2.83 | 0.63 | 0.81 | 0.73 |
| 2 | 7.66 | 2.57 | 0.59 | 0.78 | 0.70 |
| 3 | 7.63 | 2.54 | 0.60 | 0.79 | 0.71 |
| 4 | 7.49 | 3.36 | 0.64 | 0.82 | 0.74 |
| 5 | 7.41 | 2.58 | 0.67 | 0.85 | 0.76 |
| 6 | 7.40 | 2.02 | 0.68 | 0.85 | 0.77 |
| 7 | 7.40 | 2.48 | 0.77 | 0.91 | 0.83 |
| 8 | 7.19 | 2.65 | 0.66 | 0.84 | 0.76 |
| 9 | 7.14 | 2.51 | 0.61 | 0.80 | 0.72 |
| 10 | 7.14 | 2.40 | 0.54 | 0.73 | 0.67 |
| 11 | 7.09 | 2.57 | 0.63 | 0.82 | 0.73 |
| 12 | 7.03 | 2.56 | 0.67 | 0.85 | 0.77 |
| 13 | 6.99 | 2.71 | 0.67 | 0.85 | 0.76 |
| 14 | 6.98 | 1.99 | 0.64 | 0.83 | 0.74 |
| 15 | 6.97 | 2.61 | 0.40 | 0.60 | 0.55 |
| 16 | 6.90 | 2.01 | 0.61 | 0.79 | 0.72 |
| 17 | 6.82 | 2.44 | 0.61 | 0.80 | 0.72 |
| 18 | 6.79 | 2.72 | 0.65 | 0.83 | 0.75 |
| 19 | 6.76 | 2.42 | 0.70 | 0.87 | 0.78 |

The JAK kinase family consists of at least four tyrosine kinases including Tyk2, JAK1, JAK2 and JAK3, which share significant structural homology with each other^42^. To identify which one was the main target of NOT, we measured the IC50 values for each JAK kinase in a cell-free experiment, the same method used in Figure 3h. The IC50 of JAK2 and JAK3 was much lower than the other 2 members (Fig. 5a). Next, NOT was conjugated with biotin and used for the pull-down assay (Fig. 5b to d). We first confirmed that the biotin-conjugated NOT still exhibited anti-inflammation effect as the unconjugated NOT, which reduced expression of CXCL10, CXCL12, CCL5 or CXCL9 to comparable levels in TNFα-treated macrophages (Supplementary Fig. 4a), as well as expression of IL-1β, IL-6, TNFα, or iNOS2 in LPS/IFNγ-treated macrophages (Supplementary Fig. 4b). NOT-biotin could markedly pull-down JAK2 and JAK3 from the cell lysates of 293T cells that were overexpressed with the four JAK kinase family members, respectively (Fig. 5c), but showing no interaction with JAK1 or Tyk2. We also confirmed that NOT-biotin pulled down JAK2 using 293T overexpression system and PEMs (Supplementary Fig. 4c,d). Furthermore, we performed a competition assay using cell lysates from PEMs and found that in the presence of NOT, NOT-biotin pulled down less amount of JAK2 (Fig. 5d). Importantly, after we incubated NOT-biotin in the cell culture medium with macrophages followed by a pull down assay using NOT-biotin, we found that NOT-biotin could enter inside macrophages and interact with the endogenous JAK2 (Fig. 5e).

**Fig. 5.**
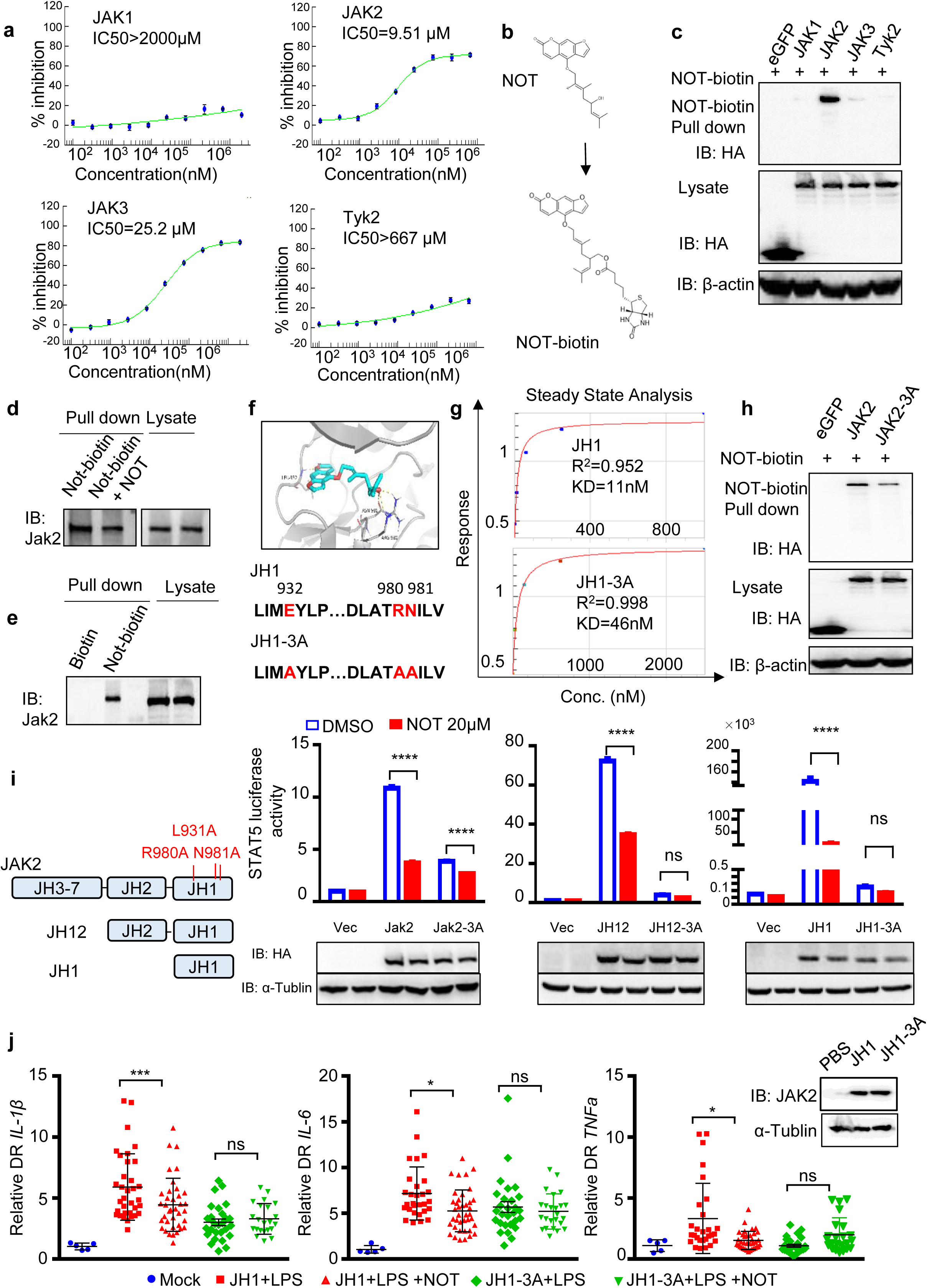
NOT inhibits JAK2 activity via directly binding to JAK2. **a**, Z’-Lyte assay was used to test the effect of NOT to inhibit the activation of JAK1/2/3 and Tyk2. **b**, The chemical structures of NOT and the biotin-conjugated NOT analog. **c,** 293T cells were transfected o express eGFP, JAK1/2/3 or Tyk2 and cell lysates were incubated with NOT-biotin, followed by pull-down assay using streptavidin beads. Representative blot of three independent experiments was shown. **d-e,** The mixture of NOT and NOT-biotin (1:1) for the competition assay, or the NOT-biotin alone, or the biotin as a negative control, were incubated with lysates of PEMs, pulled down by streptavidin beads followed by immunoblotting using anti-JAK2 antibodies. Representative blot was shown. **f,** The computer-assisted analysis of NOT binding to JAK2. NOT (blue and red) was docked in the JH1 domain (PDB ID: 4E6Q) (upper panel). The lower panel indicates the three key residues in the JH1 (808–1129) domain of JAK2 and the JH1-3A mutant (L932A/R980A/N981A). **g**, SPR analysis of NOT-biotin binding to the JH1 domain or the JH1-3A mutant by ForteBio Octet. **h**, 293T cells were transfected to express eGFP, JAK2 or JAK2-3A, and cell lysates were incubated with NOT-biotin followed by pull-down assay using streptavidin beads. **i**, 293T cells were transfected with the vector, JAK2, JAK2-3A or the vector, JH12, JH12-3A or the vector, JH1, JH1-3A plasmids (expression levels shown in lower panels) together with STAT5-luciferase plasmid, then cultured with or without NOT (20 µM) to measure the luciferase readings (Upper panels). Data represent mean ± SEM of at least three experiments. **j**, The mRNA of JH1 or JH1-3A (100 pg/1nL) were injected into one cell stage of zebrafish embryos, followed by injection of LPS (2mg/ml) with or without NOT (40 μM) at 48 hrs post-fertilization (hpf). The mRNA levels of zebrafish IL-1β, IL-6 and TNFα were checked at 54 hpf. The protein expression levels of JH1 or JH1-3A at 50 hours post-fertilization in zebrafish were confirmed by immunoblotting (right panel).

The JAK2 protein consists of a tyrosine kinase domain (JH1), a pseudo-kinase domain (JH2), an SH2 domain and a FERM domain ^43, 44^. In order to investigate which site(s) in JAK2 could bind to NOT, we used the SYBYL software (version 2.1) to mimic the possible binds sites that suggested the interaction at the R980, N981 and L938 sites (Fig. 5f). Interestingly, these sites were all located in the JH1 domain. We next mutated the three sites into alanine in the JH1 domain of JAK2 (L938A/R980A/N981A, named as JH1-3A), which were purified from *E.coli* (Supplementary Fig. 4e). The SPR analysis revealed that the affinity of NOT binding to the wild type JH1 domain was 11 nM [KD (equilibrium dissociation constant) value], which was about 5 times lower than the affinity of NOT binding to the JH1-3A mutant (46nM) (Fig. 5g). We also mutated the three sites into alanine in the full length of JAK2 (named as JAK2-3A). Consistently, the JAK2-3A mutant were pulled down by NOT-biotin at much lower levels compared to that in the WT JAK2 (Fig. 5h).

To further confirm this, 293T cells were transfected with the STAT3 or STAT5 luciferase reporter plasmids with JAK2 or JAK2-3A. Overexpression of JAK2 enhanced the STAT3 or STAT5 luciferase readings, which were significantly suppressed by NOT treatment (Fig. 5i left panel and Fig. 5j). By contrast, overexpression of the JAK2-3A mutant could not upregulate STAT5 activity(Fig. 5i right panel), which failed to be further markedly inhibited by NOT treatment (Fig. 5i left panel). To confirm the importance of the three sites in JAK2, we next overexpressed the JH1 and JH2 domain (named as JH12), or the JH1 domain or the 3A mutants, respectively with the STAT5 luciferase reporter. NOT treatment significantly decreased the JH12 or JH1 domain-induced STAT5 activity (Fig.5i middle and right panels); In contrast, the JH12-3A or JH1-3A mutants diminished STAT5 activity which could not be further reduced by NOT treatment (Fig.5i middle and right panels).

In addition, we injected mRNAs of JH1 and the JH1-3A mutant into the one cell stage embryos of zebrafish, in order to investigate how the L938A/R980A/N981A mutations in JAK2 could affect the *in vivo* inflammatory response after LPS injection. Overexpression of JH1 and JH1-3A was confirmed in zebrafish (Fig. 5k right panel), and the WT JH1 enhanced production of IL-1β, IL-6 and TNFα compared to those of the JH1-3A mutant (Fig.5, Red vs Green square). NOT treatment decreased the IL-1β, IL-6 and TNFα expression in the JH1 group, but showed no effect in the JH1-3A mutant samples (Fig. 5k Red vs Green triangle). Together, we have elucidated that NOT binds directly to JAK2 to inhibit JAK2 activation via the critical 3 sites, including R980, N981 and L938.

### Combination of NOT with anti-TNFα enhance therapeutic efficacy on CIA mice

The enhanced levels of TNFα are found in RA patients and in RA mouse models, and treatment with anti-TNFα monoclonal antibodies (e.g. etanercept and infliximab) are effective in the clinic^45, 46^. In addition, targeting the JAK family has been reported to be potential therapy for treating RA ^47, 48, 49^. Thus, we asked whether the combination of anti-TNFα and NOT could achieve a better therapeutic effect than anti-TNFα alone in the CIA model.

We treated the CIA mice with etanercept (ETA) that received twice immunization using collagen, and observed that the effect of NOT was comparable to ETA in attenuating the scores of CIA (Fig. 6a,b, Green vs Purple dots). Notably, the combination of NOT and anti-TNFα treatment further reduced the clinical scores of the CIA mice (Fig. 6a,b, Black dots). Moreover, radiographic and histomorphometry examinations revealed that the combination of NOT with anti-TNFα treatment achieved better therapeutic effects to protect mice from bone erosions, compared to NOT treatment alone or anti-TNFα treatment alone (Fig. 6c,d). This indicates that NOT could be used together with anti-TNFα to treat RA patients.

**Fig. 6.**
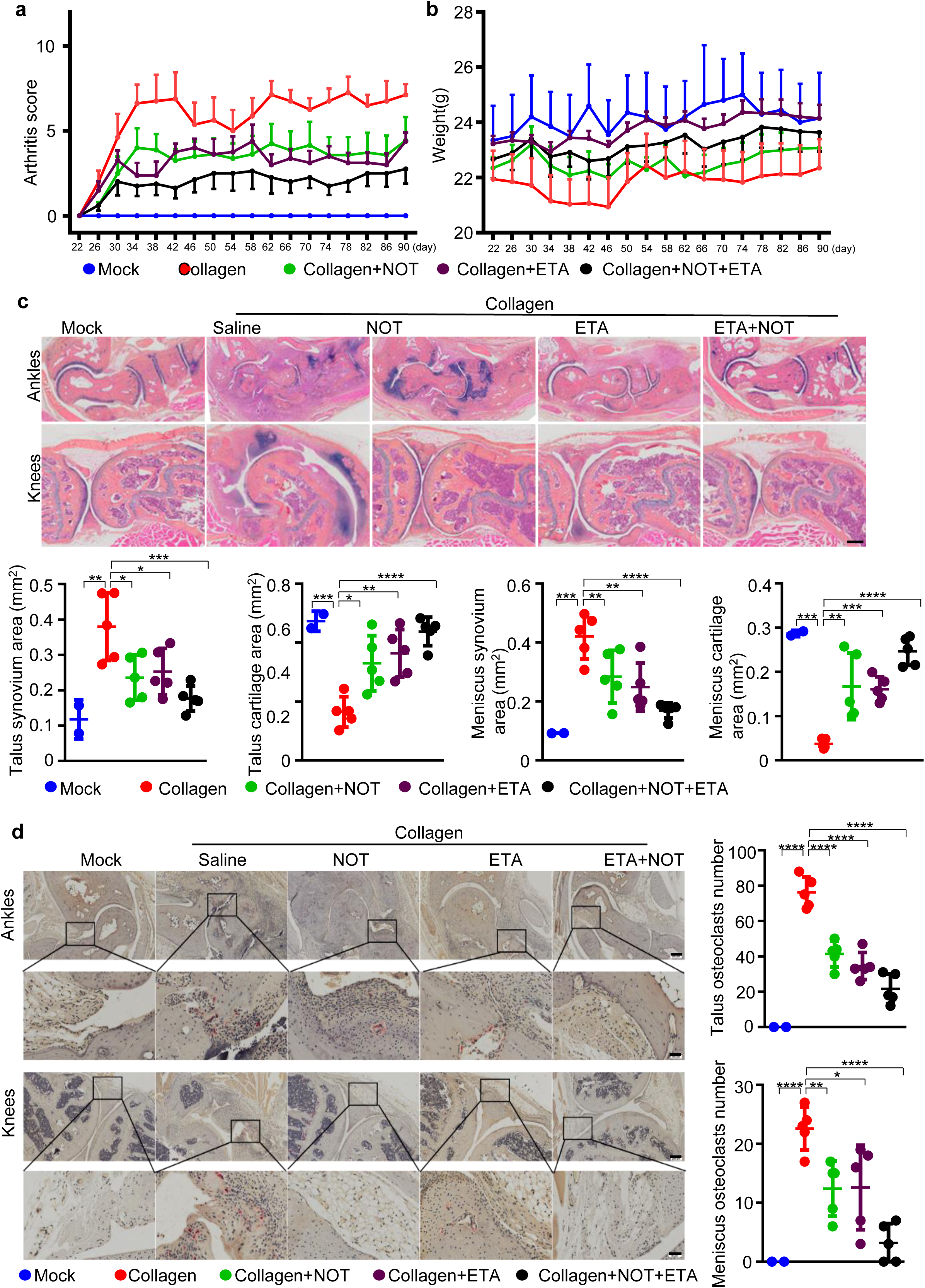
Combination of NOT with anti-TNFα (etanercept, ETA) enhance therapeutic efficacy on CIA mice. **a-b**, DBA/1J mice were twice immunized with collagen, and treated with NOT (10mg/kg), anti-TNFα (10 mg/kg), NOT (10mg/kg)+anti-TNFα (10 mg/kg) or Saline (n=8 each group) to record clinical score. Data represent mean ± SEM, two-way ANOVA. **c**, Representative photomicrographs of ABOG-stained tissue sections of ankle and knee joints (Scale bar: 100 μm). Quantification analysis of synovitis and cartilage area around talus and meniscus. Data represent mean ± SD (n=5 each group) from at least two experiments, one way ANOVA. **d**, Representative TRAP staining images showed osteoclasts in synovial tissues around talus and meniscus in the hind paw joints (Upper scale bar: 200 μm, Lower scale bar: 50 μm; n=5 each group). Data represent mean ± SD, one way ANOVA. *P<0.05, **P<0.01, ***P<0.001, ****P<0.0001.

## Discussion

NOT is the major active ingredient of the traditional Chinese medicinal herb *Notopterygium incisum Ting ex H.T. Chang.* Here, we have demonstrated for the first time that NOT is effective to treat collagen-induced arthritis in two different mouse strains using different routs including intraperitoneal injection and oral administration.

It has been reported that the monocyte-derived infiltrating macrophages dramatically increased in the flare joints in RA^6^. The flare joints are marked by co-existence of TLR4 activation and high levels of IFNγ. When macrophages are triggered by the TLR4 agonist LPS and IFNγ, a large amount of TNFα is produced which is consistent with prior reports that macrophages are a major source of TNFα in the inflammatory joint ^2, 3^. In turn, TNFα could stimulate macrophages to produce large amounts of cytokines and chemokine, resulting in an excessive inflammation to damage joint bone and cartilage in the affected joint. We have found that NOT treatment significantly reduces clinical RA scores with less inflammatory macrophage infiltration at the joints (Fig.1,2 and Supplementary Fig.1), which also directly acts on macrophage to prevent production of proinflammatory cytokines and chemokines upon TNFα or LPS/IFNγ stimulation (Fig.3). Although CIA could mimic the pathology state of RA, there were no “perfect” animal models ^50, 51, 52^. To explore the therapeutic effects of NOT, we applied three CIA models with two different NOT administration in our study, including one-time immunized DBA/1J mice, two times immunized DBA/1J and C57/BL6 mice. We found that NOT has a good therapeutic effect in reducing the clinical score, synovial area and bone destruction in all these CIA models.

More importantly, co-treatment of NOT and an anti-TNFα antibody could achieve a better therapeutic effect in the CIA mouse model than anti-TNFα treatment alone (Fig.6). Because anti-TNFα neutralizing antibodies are expensive and the side effects are notable ^53^, it is not suitable for a long-term usage. Therefore, anti-TNFα antibody (or anti-IL1 antibody and anti-IL-6 antibody) is used in combination with a small molecule such as anti-rheumatic drugs methotrexate or leflunomide to treat RA in the clinic ^54^. Our findings indicate that NOT is potentially used in combination with anti-TNFα neutralizing antibodies to improve the therapeutic effect, which might also benefit from decreasing the dose of anti-TNFα agents to reduce its side effects. Taken together, we have demonstrated that NOT is a potential drug candidate for RA patients.

Over the past decades, the JAK family has been regarded as an attractive drug target for the treatment of RA or other diseases ^55, 56^. We are excited to report that the natural compound NOT effectively acts on JAK2/3 to specifically blocks the JAK-STAT pathway. We have revealed that the three sites at 931/arginine, 980/asparagine and 981/leucine are the key sites on JAK2 for binding NOT. When these sites were mutated, JAK2-3A binds less to NOT and decreases STAT5 activation (Fig.5i). Further studies are necessary to understand exactly how the three amino acid residues are critical for NOT binding to block JAK2 activity.

Besides its therapeutic effect on arthritis, JAK2/3 are suggested to be critical targets in many diseases such as Inflammatory Bowel Disease^57^, Psoriasis ^58^, localized hair loss ^59^, allergic dermatitis ^60^ and other autoimmune diseases^61^. As an inhibitor of JAK2/3, NOT may be an effective therapeutic for these JAK2/3-involved diseases, and such possibilities require further studies especially whether NOT mainly targets JAK2/3 in macrophages or other immune cell types in these diseases. Further, the JAK inhibitors including Tofacitinib have been reported to have side effects with respect to infection^62^, anemia or leukemia ^62^, abnormal lipid metabolism and cardiovascular disease ^63^ or cancer ^64^. Although we did not find signs of toxicity when the mice were treated with NOT, including hepatocyte, glomerulus, lung or spleen injury, further examinations should be performed to understand whether NOT could safely target JAK2/3 with less side effects.

In conclusion, we have demonstrated that NOT directly binds and targets JAK2/3 to suppress macrophage-mediated inflammation, and significantly ameliorates arthritis in the CIA mouse models without obvious side effects. NOT might be a new JAK2/3 inhibitor, which could be used in combination of anti-TNFα antibodies to treat RA, as well as other JAK-related diseases.

## Acknowledgements

The authors were grateful to Gaokeng Xiao from Guangzhou Molcalx Information & Technology Ltd for the Surflex-Dock assistance. This work was supported by research grants from the Strategic Priority Research Program of the Chinese Academy of Sciences (XDB19000000 to HW), National Natural Science Foundation (81825011 to HW, 81822050 and 81673990 to LQQ, 81330085 to SQ), The program for innovative research team of ministry of science and technology of China (2015RA4002 to WYJ), “Innovation Team” development projects (IRT1270 to WYJ), Research project of Shanghai science and Technology Commission (17401971100 to LQQ), Shanghai TCM Medical Center of Chronic Disease (2017ZZ01010 to WYJ).

## Author contributions

QW, XZ and LY performed most of the experiments, analyzed the data and participated in the preparation of the manuscript. YJZ and JX bred mice, helped with i.p. injection and prepared samples. QS, YJW and QQL provided scientific input and helped with manuscript editing. HW designed the study, and drafted and finalized the manuscript. All authors read and approved the final manuscript.

---

**Supplementary Fig. 1.**
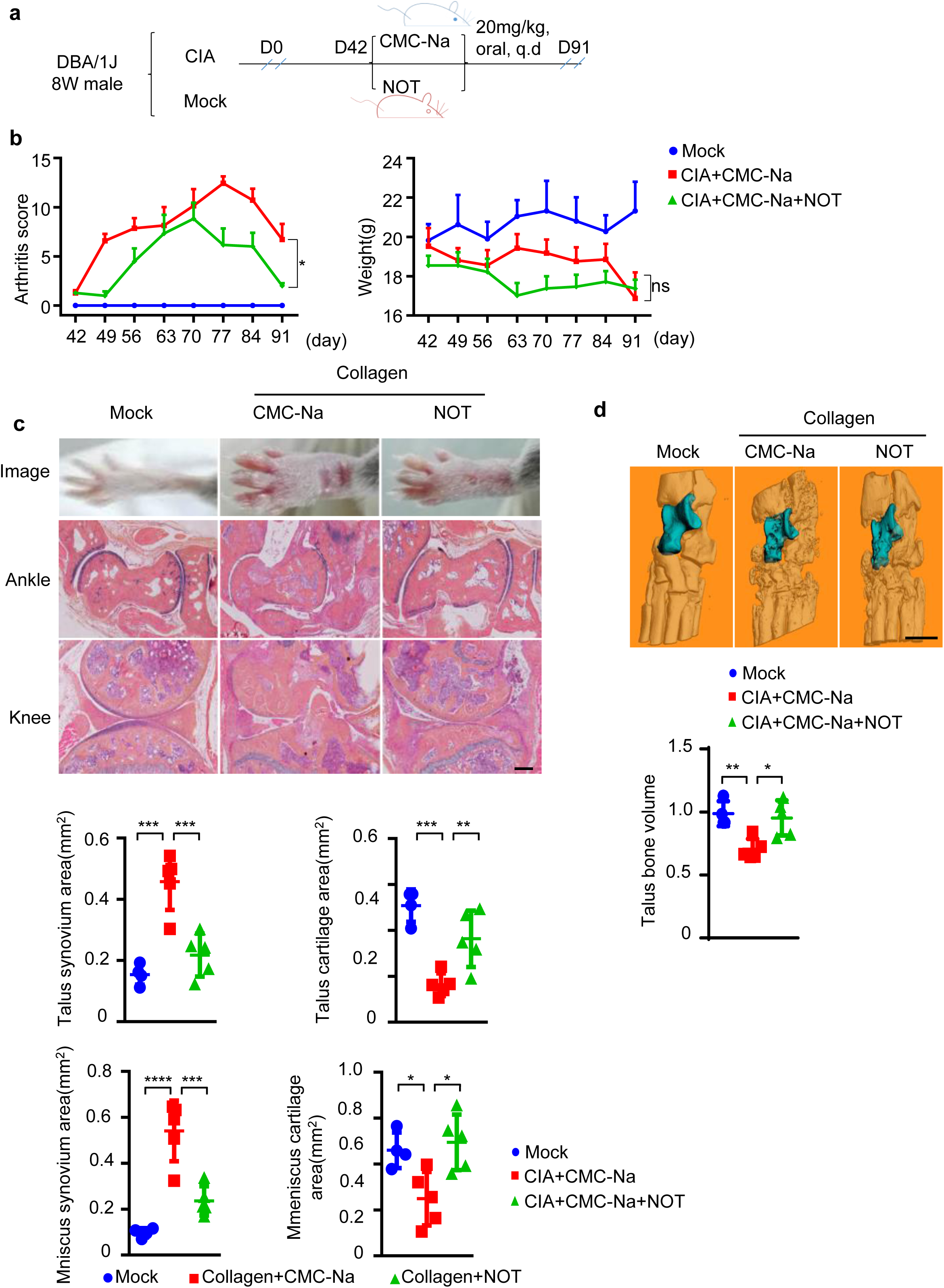
Oral administration of NOT protects DBA1/J mice from collagen-induced arthritis. **a-b**, DBA/1J CIA mice were immunized once with collagen, then intragastrically daily treated with NOT or the control CMC-Na for 49 days after first immunization (from D42 to D91). Clinical scores and body weight were recorded (mean ± SEM, n=4 Mock, n=7 CMC-Na, n=6 NOT, two-way ANOVA). **c**, Representative images of ABOG-stained tissue sections (Scale bar: 100 μm) of ankle or knee joints. Quantification analysis of synovitis and cartilage area around talus and meniscus (n=4 Mock, n=5 CMC-Na, n=5 NOT, one-way ANOVA). **d**, Representative MicroCT images (Scale bar: 1mm) and quantification of talus bone volume (n=4 Mock, n=5 CMC-Na, n=5 NOT, one-way ANOVA). Data represent mean ± SD from two experiments. *p<0.05, **p<0.01, ***p<0.001, ****p<0.0001.

**Supplementary Fig. 2.**
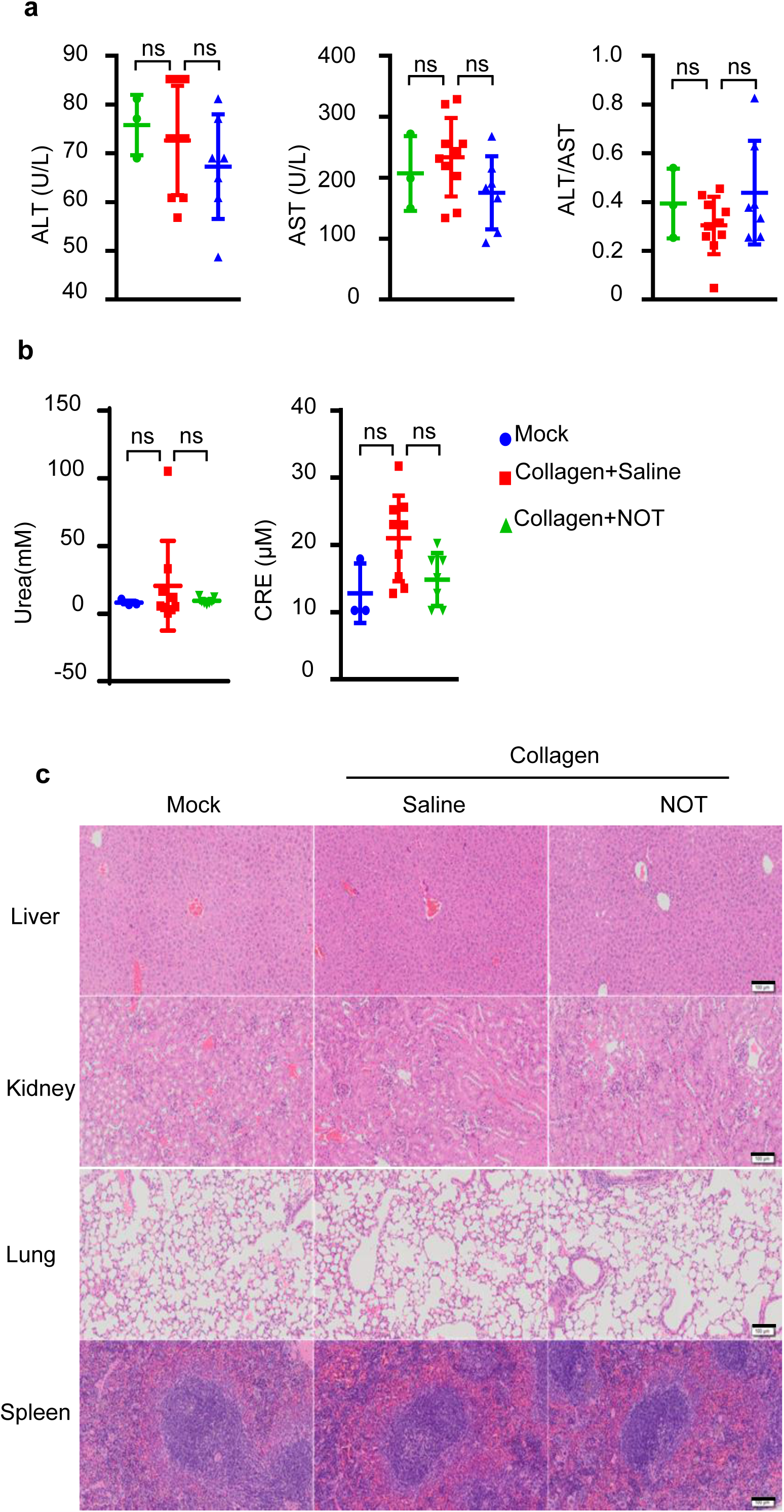
NOT did not show any damage effect on liver and kidney of CIA mice. **a**, Serum levels of aspartate aminotransferase (AST) and alanine aminotransferase (ALT) and the ratio of ALT/AST were assessed. **b**, Serum levels of creatinine (CRE) and blood urea nitrogen (UREA) were detected by the CRE reagent kit or UREA reagent kit. Data represent mean ± SD of three experiments (n=3 Mock, n=10 CIA treated with Saline, n=7 CIA treated with NOT). One-way ANOVA, ns indicates P > 0.05. **c**, Representative hematoxylin-eosin (H&E) staining of liver, kidney, lung and spleen in the Mock (n=4) or the CIA mice treated with NOT (n=10) or saline (n=7) (Scale bar: 100μm).

**Supplementary Fig. 3.**
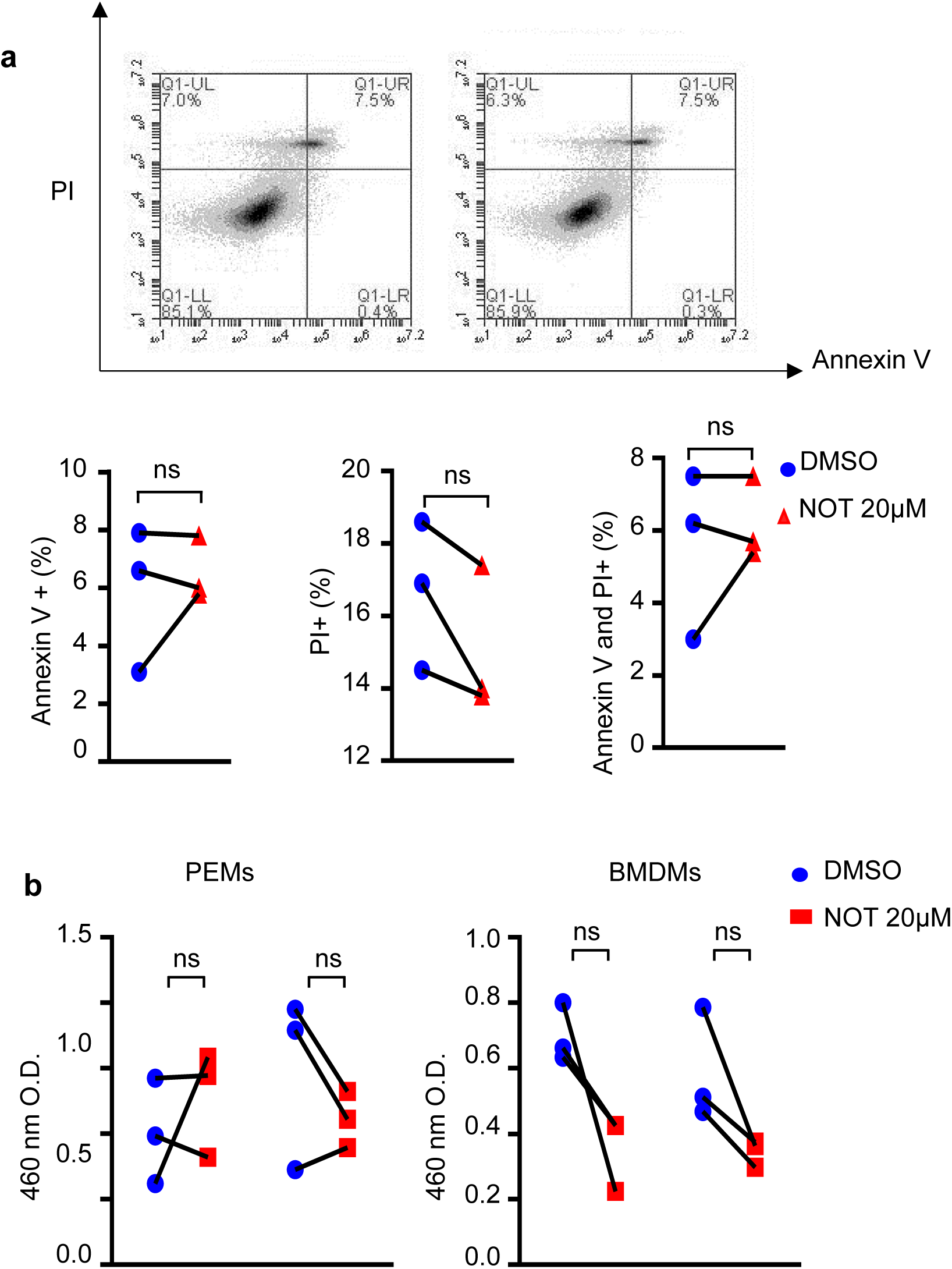
NOT does not affect macrophage apoptosis or proliferation in vitro. PEMs were treated with NOT (20 µM) or the DMSO control. **a**, Cell apoptosis was assessed by Annexin V and PI staining via flow cytometry. **b**, Cell growth was determined by the CKK8 assay. Data represent mean ± SD of four experiments. Paired Student’s t-test, ns indicates P > 0.05.

**Supplementary Fig. 4.**
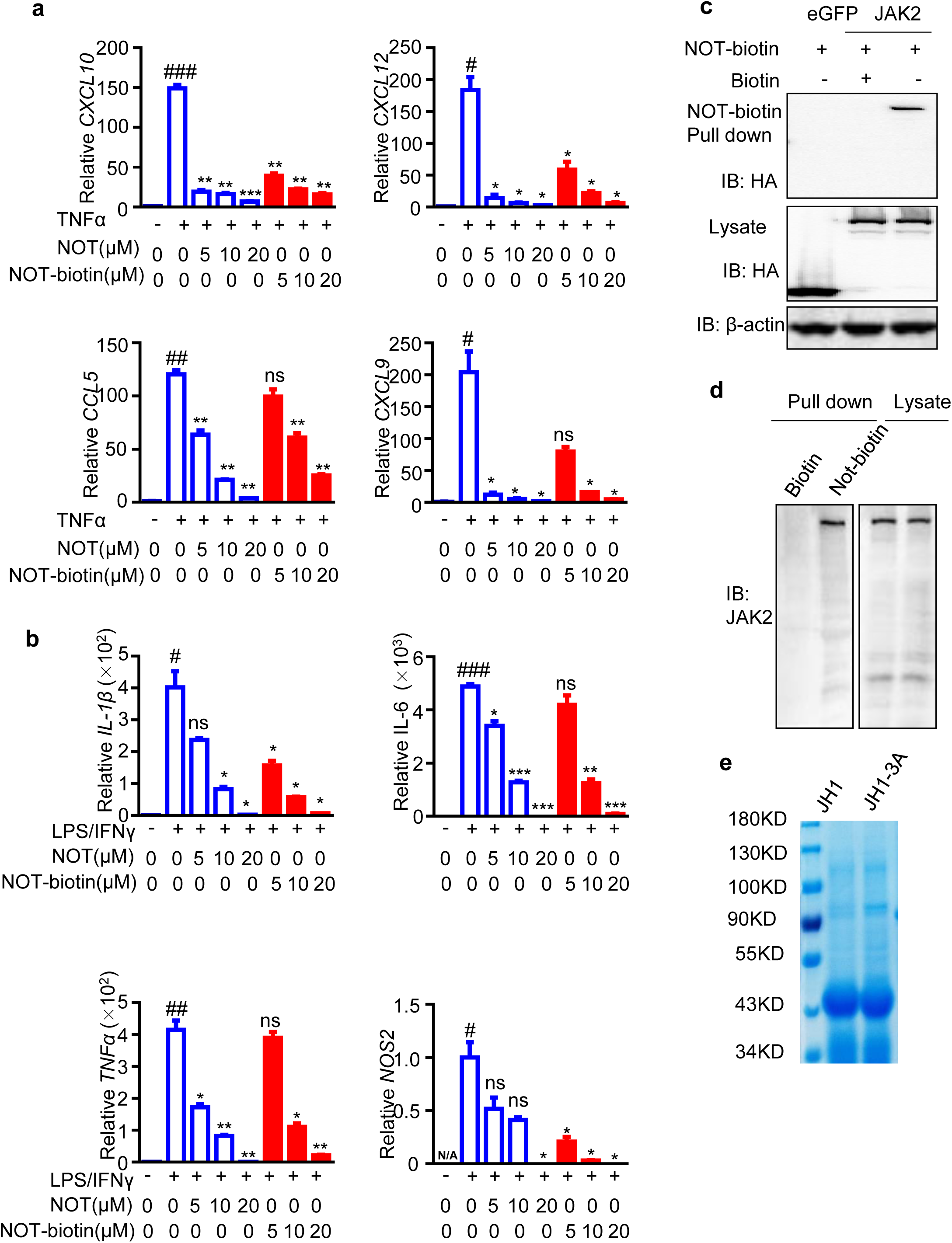
NOT inhibits JAK2 activity via binding to JAK2 directly. **a**, The mRNA levels of CXCL10, CXCL12, CCL5 and CCL9 were checked in TNFα-stimulated BMDMs, or **b**, The mRNA levels of IL-1β, IL-6, TNFα and NOS2 were checked in LPS (1μg/ml)/IFN-γ (100ng/ml)-stimulated BMDMs, which were pretreated with NOT (5, 10, and 20 μM) or NOT-biotin (5, 10, and 20 μM). Data represent mean ± SEM from at least 3 mice, one-way ANOVA. #P<0.05, ##P<0.01, ###P<0.001 were labeled when the TNFα- or LPS/IFN-γ-stimulated group were compared with the resting Mock group. And ns indicating P>0.05, *P<0.05, **P<0.01, ****P<0.0001 were labeled when the NOT-treated samples were compared with TNFα- or LPS/IFNγ-treated samples. N/A, not detected (CQ>35). **c-d**, Cell lysates of 293T cells that were transfected with the eGFP or JAK2 **(c)**, or cell lysates of PEMs **(d)** were incubated with NOT-biotin or biotin followed by pull-down assay using streptavidin beads. Representative blot is from three independent experiments. **e**, Coomassie blue staining showed the purified JH1 or JH1-3A protein from BL21 (DE3).

